# A peptide-neurotensin conjugate that crosses the blood-brain barrier induces pharmacological hypothermia associated with anticonvulsant, neuroprotective and anti-inflammatory properties following status epilepticus in mice

**DOI:** 10.1101/2024.07.15.603208

**Authors:** Lotfi Ferhat, Rabia Soussi, Maxime Masse, Grigorios Kyriatzis, Stéphane D. Girard, Fanny Gassiot, Nicolas Gaudin, Mathieu Laurencin, Anne Bernard, Angélique Bôle, Géraldine Ferracci, Maria Smirnova, François Roman, Vincent Dive, Salvatore Cisternino, Jamal Temsamani, Marion David, Pascaline Lécorché, Guillaume Jacquot, Michel Khrestchatisky

## Abstract

Preclinical and clinical studies show that mild to moderate hypothermia is neuroprotective in sudden cardiac arrest, ischemic stroke, perinatal hypoxia/ischemia, traumatic brain injury and seizures. Induction of hypothermia largely involves physical cooling therapies, which induce several clinical complications, while some molecules have shown to be efficient in pharmacologically induced hypothermia (PIH). Neurotensin (NT), a 13 amino-acid neuropeptide that regulates body temperature, interacts with various receptors to mediate its peripheral and central effects. NT induces PIH when administered intracerebrally. However, these effects are not observed if NT is administered peripherally, due to its rapid degradation and poor passage of the blood brain barrier (BBB). We conjugated NT to peptides that bind the low-density lipoprotein receptor (LDLR) to generate “vectorized” forms of NT with enhanced BBB permeability. We evaluated their effects in epileptic conditions following peripheral administration. One of these conjugates, VH-N412, displayed improved stability, binding potential to both the LDLR and NTSR-1, rodent/human cross-reactivity and improved brain distribution. In a mouse model of kainate (KA)-induced status epilepticus (SE), VH-N412 elicited rapid hypothermia associated with anticonvulsant effects, potent neuroprotection and reduced hippocampal inflammation. VH-N412 also reduced sprouting of the dentate gyrus mossy fibers and preserved learning and memory skills in the treated mice. In cultured hippocampal neurons, VH-N412 displayed temperature-independent neuroprotective properties. To the best of our knowledge, this is the first report describing the successful treatment of SE with PIH. In all, our results show that vectorized NT may elicit different neuroprotection mechanisms mediated either by hypothermia and/or by intrinsic neuroprotective properties.

## INTRODUCTION

Preclinical and clinical studies have shown that mild to moderate hypothermia is neuroprotective in situations of exacerbated neuronal death including sudden cardiac arrest with resuscitation, ischemic stroke, perinatal hypoxia/ischemia and traumatic brain injury (Kida et al., 2013; Andresen et al., 2015). Studies also suggest that hypothermia decreases seizure burden in experimental models (Sartorius and Berger, 1998; Schmitt et al., 2006; Niquet et al., 2015a,b) and in humans (Karkar et al., 2002; Kim et al., 2017). Selective brain cooling has also broad-ranging anti-inflammatory effects and prevents the development of spontaneously occurring seizures in a rat model of post-traumatic epilepsy (D’Ambrosio et al., 2013). Results in animal studies are supported by clinical data showing a positive relationship between therapeutic hypothermia (TH) and seizures in neonates with hypoxic-ischemic encephalopathy (Orbach et al., 2014). Pediatric case series report treatment of refractory status epilepticus (RSE) with mild hypothermia, which decreases seizure burden during and after pediatric RSE and may prevent RSE relapse. Hypothermia is also used in several centers around the world as second line therapy for patients with RSE, despite a small evidence base (Guilliams et al., 2013; reviewed in Ferlisi and Shorvon, 2012; Bennett et al., 2014). While the level of hypothermia is uncertain, it has been suggested that mild hypothermia is most effective (Rossetti and Lowenstein, 2011). Focal brain cooling (FBC) also reduces epileptic discharges (EDs) and concentrations of glutamate and glycerol in patients with intractable epilepsy, suggesting neuroprotective effects (Nomura et al., 2014). Current methods for the induction of hypothermia largely involve physical cooling therapies, which induce several clinical complications, including electrolyte disturbances, coagulation dysfunction, infections, cardiac arrhythmia (Carraway and Leeman, 1975, 1973). In particular, forced hypothermia lowers core temperature by overwhelming the body’s capacity to thermoregulate, but does not change the temperature set-point, thus generating counter-regulation mechanisms such as shivering and tremor (Feketa et al., 2013; Suchomelova et al., 2015), which warrant sedation and curarization in intensive care units (Andresen et al., 2015; Hammer et al., 2009). Pharmacologically-induced hypothermia (PIH) was obtained in animal models using different molecules. They promote a controlled decrease in core temperature by lowering the brain’s temperature set-point and maintaining thermoregulation at lower set points (Liska et al., 2018). Among those, neurotensin (NT) is a 13 amino-acid neuropeptide that modulates body temperature (Coquerel et al., 1988, 1986; Fanelli et al., 2015). NT interacts with 3 receptor subtypes, including NTSR1, NTSR2, and gp95/Sort-1 or NTSR3, to mediate its peripheral and central effects. The G protein-coupled receptors NTSR1 and NTSR2 have seven transmembrane domains (Vincent, 1995) while Sort1/NTSR3 only has one single transmembrane domain and is not coupled to a G protein (Mazella, 2001). The role of NT in neuroprotection and neuroinflammation, and the receptors involved, remain largely unknown. NTSR1 and NTSR2 differ in their affinity for NT, with NTSR1 and NTSR2 showing high and lower affinity respectively (Tanaka et al., 1990; Chalon et al., 1996). NTSR1 is expressed prenatally, preferentially in neurons in different brain structures (Palacios et al., 1988), while NTSR2 is expressed postnatally, essentially in glial and endothelial cells and increases during brain development (Sarret et al., 1998; Lépée-Lorgeoux et al., 1999; Yamauchi et al., 2007; Woodworth et al., 2018; Kyriatzis et al., 2021.

NT has been shown to induce PIH when administered intracerebrally (Coquerel et al., 1986, 1988; Popp et al., 2007; Fanelli et al., 2015), by inducing a downward shift of the physiological temperature set-point (Gordon et al., 2003). However, these effects are not observed if NT is administered peripherally due to its rapid processing by peptidases and poor passage of the BBB, and a number of NT analogs have been generated that are more stable than NT and that cross the BBB to induce hypothermia (reviewed in McMahon et al., 2002; Gordon et al., 2003; Orwig et al., 2009; Boules et al., 2013). These analogs have shown significant neuroprotection in several models of acute brain damage such as hypoxic ischemia, stroke and traumatic brain injury (TBI) (Choi et al., 2012; Gu et al., 2015; Zhong et al., 2020). However, to our knowledge, there are no reports on the effects of PIH in EDs. One of our main objectives was to assess such effects in experimental epileptic conditions. For this purpose, we generated “vectorized” forms of NT that cross the BBB and that display potent hypothermic properties. Indeed, transport of active principles across the BBB can be enhanced by conjugation to vector molecules designed to bind specific receptors involved in receptor-mediated transcytosis (RMT) (Pardridge, 2001, 2003; Boer & Gaillard, 2007; Jones & Shusta, 2007; Pardridge, 2007; reviewed in Vlieghe and Khrestchatisky, 2013). Several BBB receptors have been described that undergo RMT, including the transferrin receptor (TfR), the insulin receptor (IR), the insulin-like growth factor receptor (IGFR), and receptors of the low-density lipoprotein receptor (LDLR) family. A number of antibodies, protein ligands or peptides that bind some of these receptors have been developed as vectors to carry pharmacological payloads across the BBB (Friden et al., 1991; Wu et al., 1997; Wu and Pardridge, 1998; Boado et al., 2007; Pan et al., 2004; Spencer and Verma, 2007).

LDLR is part of a group of single transmembrane glycoproteins, referred to as cell surface endocytic receptors. They bind apolipoprotein complexes and are expressed with some degree of tissue specificity (Brown and Goldstein, 1979; Herz and Bock, 2002). We previously described the rational characterization and optimization of a family of cyclic peptides that bind the LDLR *in vitro* and *in vivo*. These peptides bind the EGF-precursor homology domain of the LDLR and thus do not compete with LDL binding on the ligand-binding domain. To our knowledge, they have no beneficial or untoward effects on LDL binding and LDLR activity (Malcor et al., 2012; Jacquot et al., 2016; David et al., 2018; Varini et al., 2019; Acier et al., 2021, Yang et al., 2023; Broc et al., 2024). These peptides can transport across the BBB and into the CNS and specific organs, different payloads in a LDLR-dependent manner. We and others have shown that such payloads include fluorophores, proteins, nanoparticles and liposome-based cargos (Malcor et al., 2012; Zhang et al., 2013; Chen et al., 2017; Molino et al., 2017; Cui et al., 2018; David et al., 2018; Shen et al., 2018). In the present work, we conjugated several of our LDLR-targeting peptides to NT and to shorter active variants of NT (residues 6-13 and 8-13) with different linkers. These conjugates displayed binding potential to both the LDLR and NTSR-1 receptors, with rodent/human cross-reactivity, enhanced metabolic stability in plasma compared to the native NT, and improved brain penetration potential. We selected the VH-N412 conjugate for further studies in mouse, owing to its potent hypothermia following intravenous (i.v.) administration at low dose together with optimal chemistry and conjugation features. The hypothermic and neuroprotective potential of VH-N412 was tested in the mouse model of kainate (KA)-induced status epilepticus (SE), a model of seizures associated with neurodegeneration, neuroinflammation and network reorganization. Following induction of SE, we show that the VH-N412 compound elicited rapid hypothermia that was associated with anticonvulsant effects. Seven days following SE, we observed potent neuroprotection and reduced inflammation in the hippocampus, a highly vulnerable structure to damage at early stages of epilepsy. Neuroprotection elicited by VH-N412 also reduced significantly aberrant sprouting of the dentate gyrus (DG) mossy fibers assessed 2 months after SE and preserved learning and memory skills in the treated mice. We showed that NTSR1, one of the NT receptors, was expressed in hippocampal pyramidal neurons *in vitro* and *in vivo*, in cell bodies, dendrites and spines, the post-synaptic compartment of glutamatergic synapses. Besides the neuroprotective hypothermia effects observed *in vivo* with VH-N412, we show in cultured hippocampal neurons challenged with KA and NMDA that activate glutamate receptors, that VH-N412 displayed temperature-independent neuroprotective properties that are as potent as Oestradiol /or BDNF.

## RESULTS

### Synthesis and purification of peptide-NT conjugates and their hypothermic potential in mice

We conjugated the 8-mer VH445 cyclic peptide vector that binds the LDLR (peptide 22: [cMPRLRGC]_c_, (Malcor et al., 2012) to the NT tridecapeptide through its lysine in position 6 using a three-step reaction sequence. Both peptides were prepared by solid-phase peptide synthesis on a CEM Liberty microwave peptide synthesizer using standard Fmoc/tert-butyl chemistry. Cyclization of the VH445 peptide was performed on crude peptides (Malcor et al., 2012) by formation of a disulfide bridge between the two VH445 cysteine residues. K_3_[Fe(CN)_6_] was used as an oxidating reagent. A sulfo-N-[□-maleimidocaproyloxy]succinimide ester (sulfo-EMCS) was used to incorporate a maleimido hexanoic acid linker (MHA) at lysine 6 of NT, resulting in [Lys(MHA)6]NT. In parallel, a reactive thiol moiety was incorporated on the VH445-G modified peptide by derivatization of the acid C-terminal (C-ter) with cysteamine. Conjugation was performed between both functionalized intermediates [Lys(MHA)6]NT and VH445-G-(CH_2_)_2_-SH leading to conjugate VH-N21 (Figure 1A). VH-N21 and NT were administered i.v. (bolus) in the tail vein of Swiss CD-1 mice and body temperature was monitored using digital thermometer rectal probes at different time points following administration. VH-N21 induced a transient mild hypothermia, that was dose-dependent and that reached a maximum of −3.7°C at 1 H after injection, with a dose of 8 mg/kg molar equivalent NT (eq. NT), whereas no significant hypothermia was observed with native NT at the same dose (Figure 1B). We next generated other peptide-NT conjugates based on peptides with improved properties in terms of stability and binding to the LDLR (Jacquot et al., 2016; David et al., 2018). In particular, the VH4129 peptide ([cM”Pip”RLR”Sar”C]_c_) was chosen for its optimal stability/binding properties (Jacquot et al., 2016). The plasma metabolic stability of VH4129 was shown to be higher than that of VH445 (t_1/2_ 7 H vs. 3 H, respectively), owing to a rationally optimized insertion of non-natural amino-acids. When compared with VH445, the LDLR binding of VH4129 was overall similar (K_D_ 60-70 nM). However, compared with the VH445 peptide, VH4129 presents the highest association rate (19.2 vs. 7.6 x10^5^ s^-1^.M^-1^ for VH4129 and VH445, respectively) and the highest dissociation rate (12.2 vs. 5.9 x10^-2^ s^-1^ for VH4129 and VH445, respectively). These properties provide excellent potential for peptide binding to BBB-exposed LDLR while allowing efficient release in the parenchymal compartment. Consistently, using the same initial thiol-maleimide coupling strategy as for the VH-N21 conjugate, the resulting VH-N41 conjugate (Figure 1C) induced a stronger and more sustained hypothermia after i.v. injection in mice, with a maximal body temperature decrease of −6.8°C (Figure 1D). We next evaluated a series of conjugates allowing one-pot linear synthesis with different versions of NT (residues 2-13, 6-13 and 8-13), while spacing the LDLR-targeting peptide from the neurotensin peptide using linkers such as the glycine tripeptide (GGG), aminohexanoic acid, (Ahx) or polyethylene glycol (PEG6). In this strategy, full size conjugates were synthesized in a one-step procedure on a CEM Liberty microwave peptide synthesizer using standard Fmoc/tert-butyl chemistry except for conjugates containing a PEG6 linker that was introduced manually. Cyclization was performed on crude peptides with the same procedure as described for peptide-vectors. With this new strategy synthesis yields were significantly increased compared to our initial thiol-maleimide conjugation strategy (Table S1). The conjugates were all evaluated for their potential to induce hypothermia in mice (Table S2) and allowed the selection for further studies of the VH-N412 conjugate that encompasses a PEG6 linker between the VH4129 peptide and NT(8-13) (Figure 1E). Importantly, this VH-N412 conjugate elicited a similar hypothermic response than VH-N41, with a maximal body temperature decrease of −6.4°C, but with an even more sustained profile (Figure 1F). No effect was observed with the control PEG6-NT(8-13) compound, confirming the involvement of the VH4129 peptide in the hypothermic effect of VH-N412. Dose-response curves confirmed that VH-N412 displayed an ED50 similar to that of VH-N41, estimated at 0.69 and 0.93 eq. NT, respectively (corresponding to 0.80 and 1.67 mg/kg, respectively) (Figure 1G). With its smaller size and easier production using one-pot linear synthesis, thereby leading to higher synthesis yields, VH-N412 was selected for further investigation in the mouse model of KA-induced SE.

**Figure 1:**
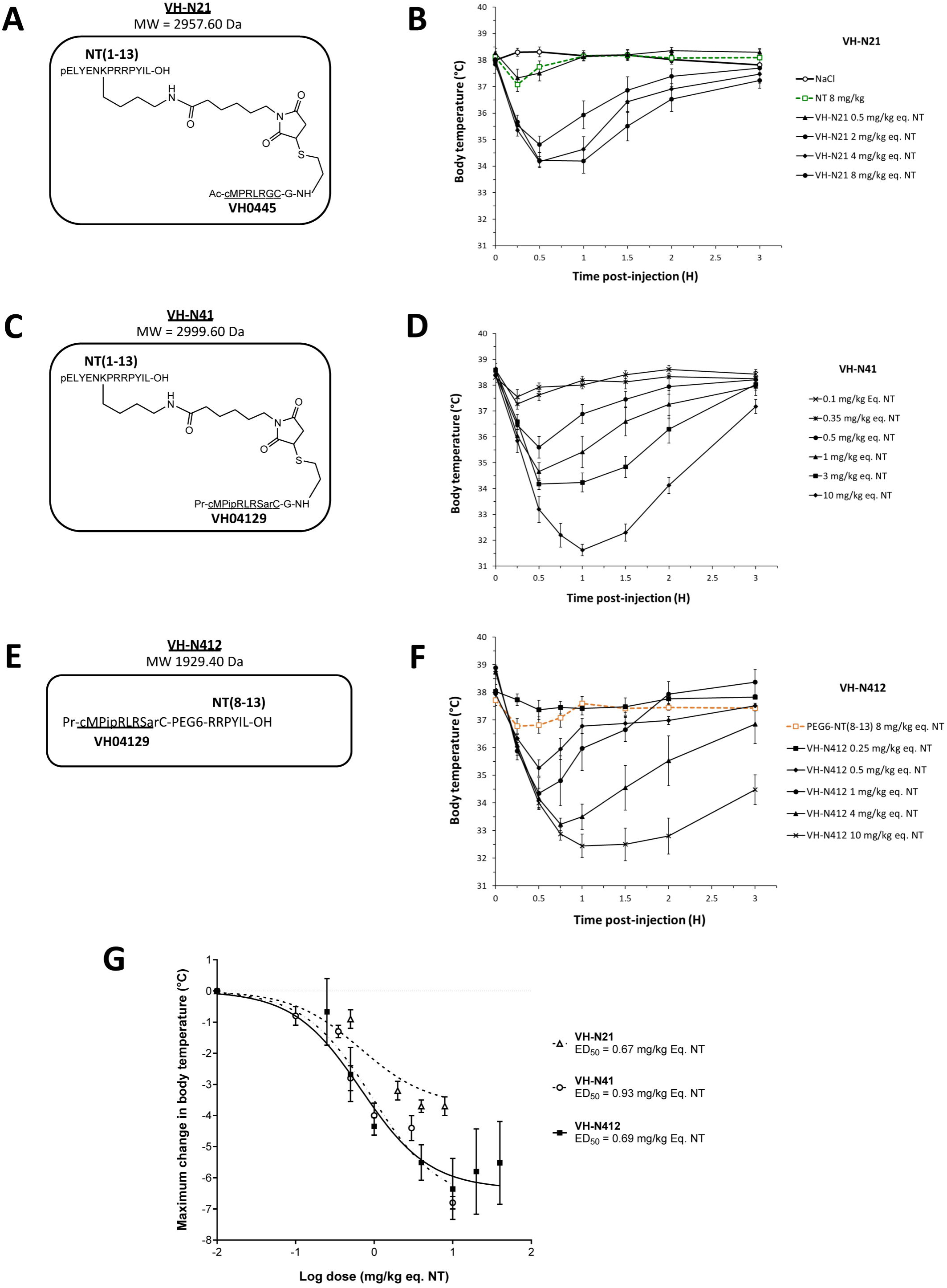
Hypothermic dose-response relationship of different VH-NT conjugates following single IV (bolus) injection in naïve Swiss (CD-1) mice. A), C) and E) show chemical structure and molecular weight of the VH-N21, VH-N41 and VH-N412 conjugates, containing the 8 amino-acid cyclic VH brain penetrating peptide that recognizes the LDL-receptor (VH445 for VH-N21 and VH4129 for VH-N41 and VH-N412), and either the NT tridecapeptide (VH-N21 and VH-N41) or it C-terminal NT(8-13) fragment (VH-N412). B), D) and F) Hypothermic response to VH-21, VH-N41 and VH-N412 conjugates in mice after single IV (bolus) injection at increasing dose-levels. Core body (rectal) temperature was measured before (baseline) and at indicated times after injection. Data are presented as means ± SEM., n = 4-8 per group. G) Dose-response curves of VH-21, VH-N41 and VH-N412 hypothermic response. ED_50_ values for each conjugate were estimated by plotting the response vs. log[dose(mg/kg eq. NT)] followed by nonlinear regression (three parameters) using GraphPad PRISM software.

### *In vitro* biological properties of VH-N412: LDLR- and NTSR1-binding, plasma stability, and *in situ* mouse brain perfusion

In parallel with the *in vivo* selection of VH-N412, we verified *in vitro* that conjugation of its VH4129 peptide and NT(8-13) moieties does not interfere with its potential to bind the LDLR. This was confirmed using Surface Plasmon Resonance (SPR) on immobilized human LDLR, with free VH4129 and the VH-N412 conjugate displaying very similar binding affinity and profiles, namely 72.6 nM and 63.8 nM, respectively (Figure 2A). We also studied the binding properties of the VH-N412 conjugate to NTSR-1. Both native NT(1-13) and VH-N412 were assessed for binding competition with a reference radiolabeled NT on cell membrane extracts expressing either the rat or the human form of NTSR-1. Both NT(1-13) and VH-N412 showed similar Ki values for rNTSR-1 and hNTSR-1, in the low nanomolar range (Figure 2B, right and left graphs). Next, the proteolytic resistance of VH-N412 was evaluated and compared to the native NT as well as the initial VH-N21 conjugate by incubation in freshly collected mouse blood at 37°C followed by quantification of the parent compound in the plasma fraction using LC/MS-MS. As opposed to the very low resistance of the native NT (t_1/2_ of 9 min), both VH-NT conjugates showed greatly enhanced stability, with VH-N412 showing the highest *in vitro* half-life estimated at 83 min, compared to 44 min with VH-N21 (Figure 2C). The blood *in vitro* half-life of VH-N412 was estimated at 74 min in human blood, demonstrating similar metabolic resistance across species (Figure 2D).

**Figure 2:**
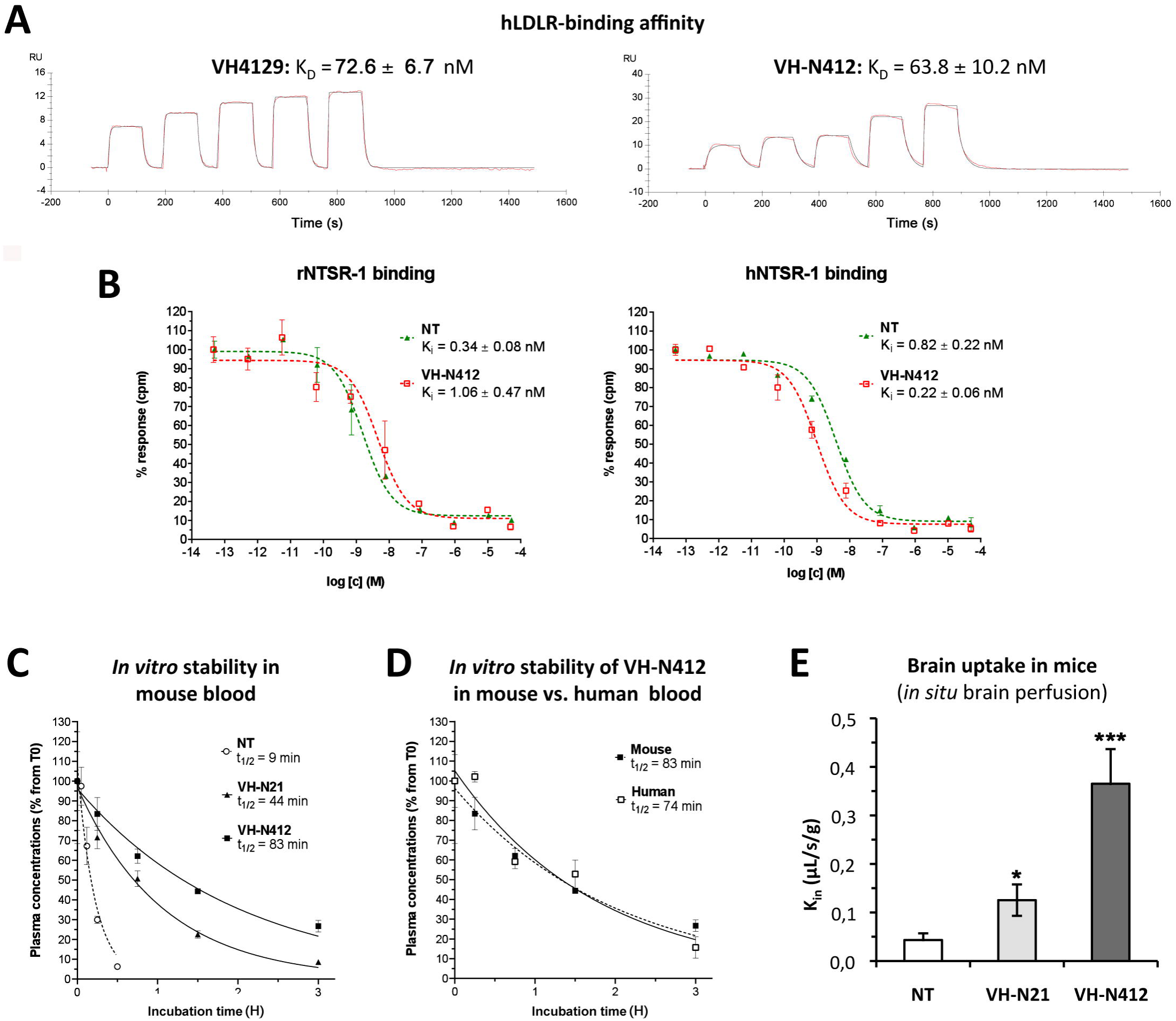
*In vitro* biological properties of VH-N412. A) Surface plasmon resonance (SPR) sensorgrams of the free VH4129 and the VH-N412 compound on immobilized hLDLR. Red lines show the specific binding of molecules obtained after double subtraction of the signal measured on the control flow cell (without immobilized LDLR) and a blank run. Black lines show fit curves of the experimental data with a 1:1 binding model. The illustrated data are representative of 2-5 independent experiments. B) shows dose-response inhibition curves of tritiated NT, bound on hNTSR-1 or rNTSR-1 membrane extracts, in the presence of indicated concentrations of NT or VH-N412. Indicated Ki values were estimated from mean IC50 values obtained by logarithmic regression of experimental data. Data plotted are means ± SD of biological duplicates. C) and D) Comparison of degradation rates for NT or VH-NT conjugates in mouse blood. NT or peptide-NT conjugates were incubated in freshly collected mouse (C) or human (D) blood and analyzed using LC-MS/MS at indicated times in the plasma fraction. Data are plotted as means ± SD of n=3 biological replicates. T_1/2_ values were estimated from nonlinear regression (one-phase-decay) of experimental data. E) BBB transport of tritium labelled NT or VH-NT conjugates using *in situ* brain perfusion in mice. Data were presented as mean ± SEM for 3 to 6 animals. Student’s *t*-test vs. NT: * *p*<0.05, ** *p*<0.01.

The mouse *in situ* brain perfusion method described by Dagenais et al., (2000) was used to measure BBB transport rate clearance (K_in_) of tritium-labelled NT, VH-N21 and VH-N412 conjugates. Consistent with the previously reported NT K_in_ of 0.013 µL.s^-1^.g^-1^ measured in mice (Gevaert et al., 2016), NT demonstrated a very low BBB transport, with a K_in_ of ∼0.04 µL.s^-1^.g^-1^ of brain tissue (Figure 2E). In contrast, VH-N21 and VH-N412 showed K_in_ values of 0.13 and 0.37 µL.s^-1^.g^-1^ respectively (Figure 2E), demonstrating that conjugating NT with these peptide-vectors enhanced its BBB transport. Furthermore, VH-N412 did not alter the integrity of the BBB. Indeed, the brain distribution volume of ^14^C-sucrose as a marker of brain vascular volume in VH-N412 mice (19.00 ± 1.00 µL.g^-1^), was in the normal range (i.e. Vvasc < 20.00 µL.g^-1^) (Cattelotte et al., 2008), and similar to that of NT (18.00 ± 2.00 µL.g^-1^).

Taken together, these results demonstrate that the VH-N412 conjugate retains its binding potential to both the LDLR and NTSR-1 receptors, with rodent/human cross-reactivity. VH-N412 encompasses the VH04129, peptide vector with higher association and dissociation rates to LDLR compared to VH0445. VH-N412 displayed greatly enhanced metabolic stability in plasma compared to the native NT, but also to the initial conjugate VH-N21, and displayed higher Kin properties with sharp improvement of brain penetration potential compared to VH-N21. These combined features contribute to the high hypothermic potential of VH-N412, requiring plasma resistance, improved BBB permeability and potent binding to its pharmacological target, namely brain NTSR1, making it an ideal candidate for further investigation of its central pharmacological potential in pathophysiological situations *in vivo*. Finally, tolerability studies were performed in naïve mice with the administration of up to 20 and 40 mg/kg eq. NT (i.e. 25.8 and 51.6 mg/kg of VH-N412) with n=3 for these doses. The rectal temperature of the animals did not fall below 32.5 to 33.2°C, similar to the temperature induced with the 4 mg/kg eq. NT dose. We observed no mortality or notable clinical signs other than those associated with the rapid HT effect such as a decrease in locomotor activity. We thus report a very interesting therapeutic index since the maximal tolerated dose (MTD) was > 40 mg/kg eq. NT, while the maximum effect is observed at a 10x lower dose of 4 mg/kg eq. NT and an ED50 established at 0.69 mg/kg as shown in Figure 1G. Severe hypothermia could also be induced in rats with different conjugates similar to VH-N412 with the same efficacy and safety (data not shown).

### Effect of VH-N412 in a model of KA-induced seizures

We assessed our VH-N412 conjugate in a model of KA-induced seizures using adult male FVB/N mice. This mouse strain was selected as a reliable and well described mouse model of epilepsy, where seizures are associated with cell death and neuroinflammation (Schauwecker, 2003; Wu et al., 2021). KA was administered subcutaneously (s.c.) in FVB/N mice at the dose of 45 mg/kg. The scheme in Figure 3A shows the timeline of the experiments we performed including physiological, histopathological, behavioral and synaptogenesis assessment. Five groups of mice were generated: SHAM, SE, SE + VH-N412, SE + NT(8-13), SE + diazepam (DZP). Body temperature was monitored before KA injection and every 30 min during 2.5 H thereafter, using a rectal probe. SE occurred around 2 H after KA injection (KA-2H) and was characterized by stage 5-6 seizures and often associated with some hyperthermia (non-significant, Figure 3B; Table S3). VH-N412 administered at the dose of 4 mg/kg eq. NT 30 min after SE onset (SE30), hence 2.5 H after s.c administration of KA (KA-2H), invariably led to transient hypothermia (Figure 3B; Table S3), which persisted at least 2 H. Mean decreases in body temperature of −2.12°C were recorded at SE30 for SE + VH-N412 animals (36.50 ± 0.34°C, *p*<0.01, Tukey’s test) as compared to SE animals (38.00 ± 0.37°C) (Figure 3B; Table S3). This hypothermia was associated with a significant decrease of seizures in the SE + VH-N412 group (1.97 ± 0.36, *p*<0.01, Tukey’s test) at SE30 as compared with the SE group (5.38 ± 0.15) (Figure 3C; Table S4). SE + VH-N412 animals presented an average of seizure intensity score of 2 or less during the rest of the experiment (SE30-SE150 min). A subset of animals was administered i.p. a high dose of DZP (15 mg/Kg), used as a positive control for its anticonvulsant effects in seizure models (for review see Sharma et al., 2018) and its hypothermic effects (Vinkers et al., 2009). As for SE + VH-N412 animals (average seizure intensity score > 5 at SE30), SE + DZP animals rapidly showed an average seizure intensity score of 1-2 during the rest of experiments, and significant hypothermia was also observed in these animals at all time points (Figure 3B,C; Tables S3 and S4; *p*<0.01, Tukey’s test). No significant variations of body temperature or seizure intensity score were observed when NT8-13 was administered at SE30, as compared with SE animals at all time points (Figure 3B,C; Tables S3 and S4).

**Figure 3:**
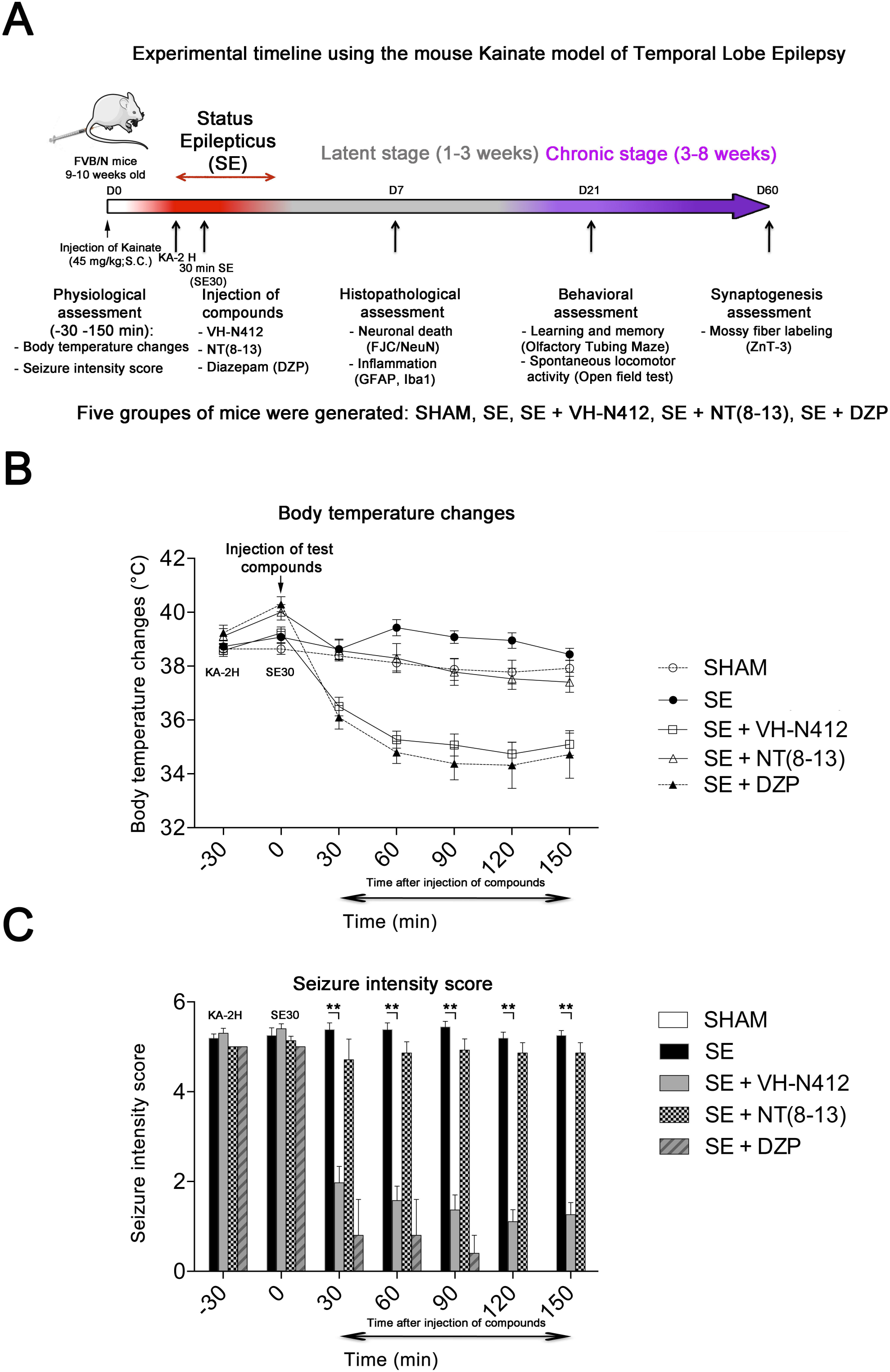
Effects of VH-N412 on body temperature and seizure intensity following status epilepticus (SE). A) shows the experimental timeline with the assessment of physiological, histopathological, behavioral and synaptogenesis features associated with the mouse KA model of temporal lobe epilepsy. Five groups of mice were generated: SHAM, SE, SE + VH-N412, SE + NT(8-13), SE + DZP. B) Mice were injected with KA, which induced stage 5 or stage 6 seizures after 2 H, characteristic of SE, associated with hyperthermia as compared to all animal groups. VH-N412 was administered 30 min after SE onset at the dose of 4 mg/kg eq. NT caused significant hypothermia, which persisted at least 2 H, similar to the effects of high dose DZP (45 mg/kg) administered i.p and used as positive control. SE + NT(8-13) had no effect on body temperature when administered 30 min after SE onset. C) Hypothermia induced by VH-N412 was associated with a significant decrease of seizures in the SE + VH-N412 group, similar to DZP, while SE + NT(8-13) had no effect on seizure intensity.

### Effects of VH-N412 on neurodegeneration and inflammation in hippocampus

Rodents that experienced SE developed inflammation and lesions in several brain areas. In our study, we focused on the hippocampal formation, where inflammation and cell death occur within the first days following KA-induced SE (Gröticke et al., 2008; Lévesque & Avoli, 2013; Li and Liu, 2019).

#### Effects of VH-N412 on hippocampal neurodegeneration

The effects of VH-N412 on hippocampal neural cell degeneration were assessed 7 days after SE (Figure 4A) using Fluoro-Jade C (FJC) staining in SHAM, SE, SE + VH-N412, SE + NT(8-13), SE + DZP animals (Figure 4A). Representative photomicrographs of hippocampal pyramidal cells from Cornu Ammonis areas 1 (CA1), 3 (CA3), Hilus (H) and granule cell layer (GCL) areas that were quantified are shown in Figure 4B while high magnification of these same areas stained with FJC in all groups of animals are shown in Figure 5. Semi-quantitative analysis revealed that FJC staining is significantly increased in CA1 (222.73 ± 11.80%, 123%, *p*<0.01, Tukey’s test), CA3 (197.27 ± 14.58%, 97% *p*<0.01, Tukey’s test) and to a lesser extent in H (120 ± 4.98%, 20%, *p*<0.05, Tukey’s test) of SE animals compared to SHAM animals (CA1: 100 ± 2.42%; CA3: 100 ± 3.18%; H: 100 ± 4.62%). No difference in FJC staining was found in the GCL (100 ± 5.44%; n=3 mice; *p*>0.05; ANOVA) (Figure 4C). These results indicate that there is major neural cell death in all hippocampal layers, including CA1-3 pyramidal cell layers and H of the DG. Neural cell degeneration observed in SE animals, was significantly decreased when VH-N412 was administered at SE30 (CA1: 97.97 ± 2.60%; CA3: 109.58 ± 7.31%; H: 89.82 ± 3.20%; *p*<0.01, Tukey’s test) (Figure 4C). In contrast, no changes were observed when NT8-13 was administered (CA1: 222.59 ± 12.02%; CA3: 200.08 ± 11.84%; H: 100.98 ± 5.75%; *p* >0.05; ANOVA) (Figure 4C). A subset of animals was administered i.p. with a high dose of DZP (15 mg/Kg) used as a positive control for its neuroprotective effects in seizure models. The results obtained for SE + VH-N412 animals were not different from those observed in SE + DZP mice (CA1: 99.21 ± 2.88%; CA3: 106.12 ± 3.56%; H: 102.07 ± 2.79%; *p*>0.05; ANOVA) (Figure 4C).

**Figure 4:**
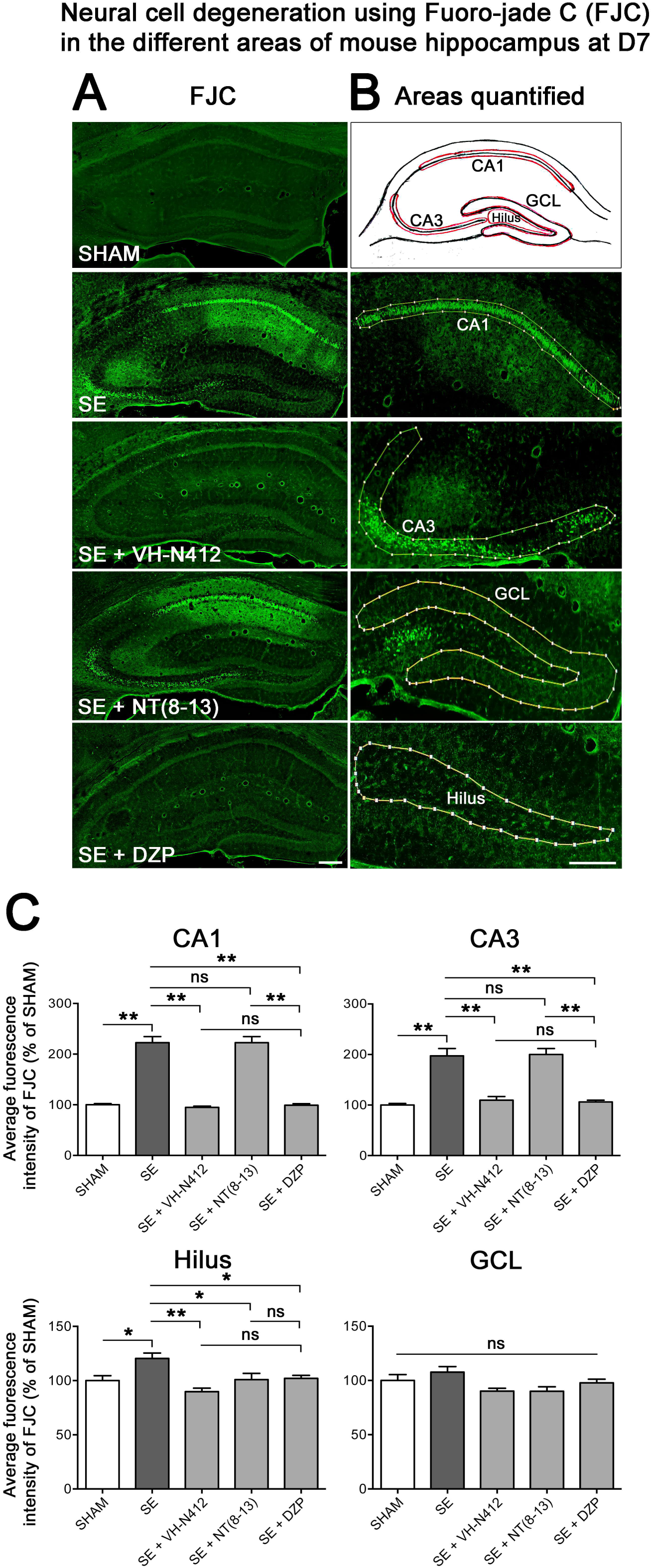
Effects of VH-N412 on neural cell degeneration following KA-induced SE. A) FJC staining was used to assess the extent of neural cell damage in coronal sections of the dorsal hippocampal formation at D7 post-SE from SHAM, SE, SE + VH-N412, SE + NT(8-13) and SE + DZP animals. B) The regions of interest are highlighted on the scheme, upper panel and were traced to quantify FJC in the CA1, CA3, GCL and the H. Scale bars: 200 μm in all panels. C) Histograms compare the mean intensities of staining for FJC in dorsal CA1, CA3, H and GCL from SHAM, SE, SE + VH-N412, SE + NT(8-13) and SE + DZP animals. VH-N412 as well as DZP displayed significant protective effect in dorsal CA1, CA3, and H but not in GCL. Data were expressed as the average percentage ± SEM. normalized to the SHAM CTL. Asterisks indicate statistically significant differences: \**p*<0.05, ** *p*<0.01 (Tukey’s-test).

**Figure 5:**
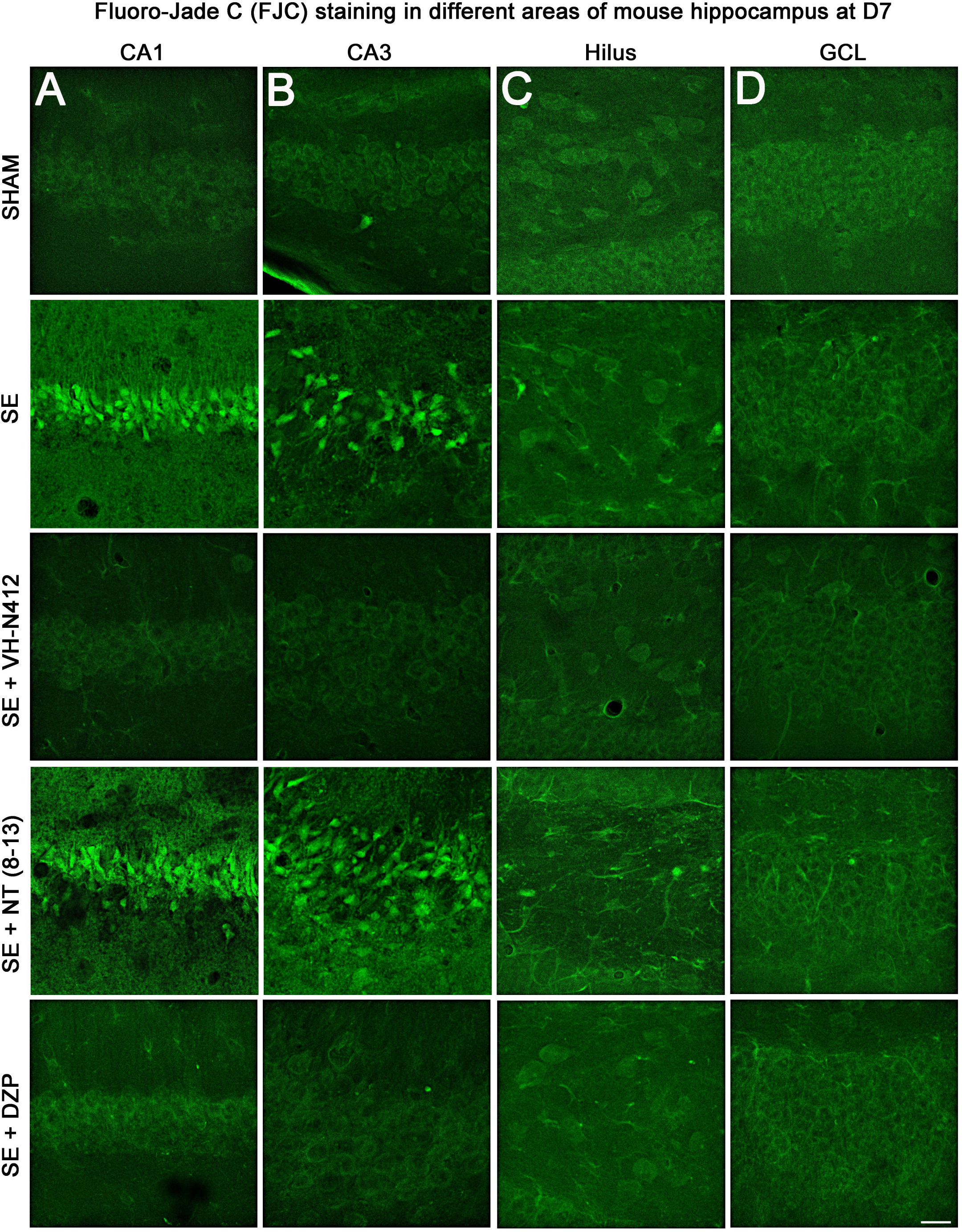
Representative examples of FJC staining in the different areas of mouse dorsal hippocampus. A) CA1, B) CA3, C) H and D), GCL, at D7 post-SE from SHAM, SE, SE + VH-N412, SE + NT(8-13) and SE + DZP animals. Scale bar: 20 μm in all panels.

Immunohistochemistry for the neuronal marker NeuN was also performed to confirm neuronal degeneration, and to evaluate on NeuN and FJC sections the effects of VH-N412 in animals at 7 days post SE. Representative photomicrographs of neurodegeneration in SE animals (compare SHAM vs SE) and neuroprotection mediated by VH-N412 are shown (Figure 6, in green, left panels). SE animals displayed a decreased NeuN staining and significantly increased neuronal death score in the hippocampal formation compared to SHAM animals. When VH-N412 or DZP was administered at SE30, the neuronal death score was significantly reduced by 51% and 34% respectively (*p*<0.01, Tukey’s test; Figure 6B, left histogram). However, no changes were observed when NT8-13 was administered (*p*>0.05; ANOVA) (Figure 6B, left histogram). Altogether, these results indicate that SE-induced neurodegeneration is partially prevented by VH-N412.

**Figure 6:**
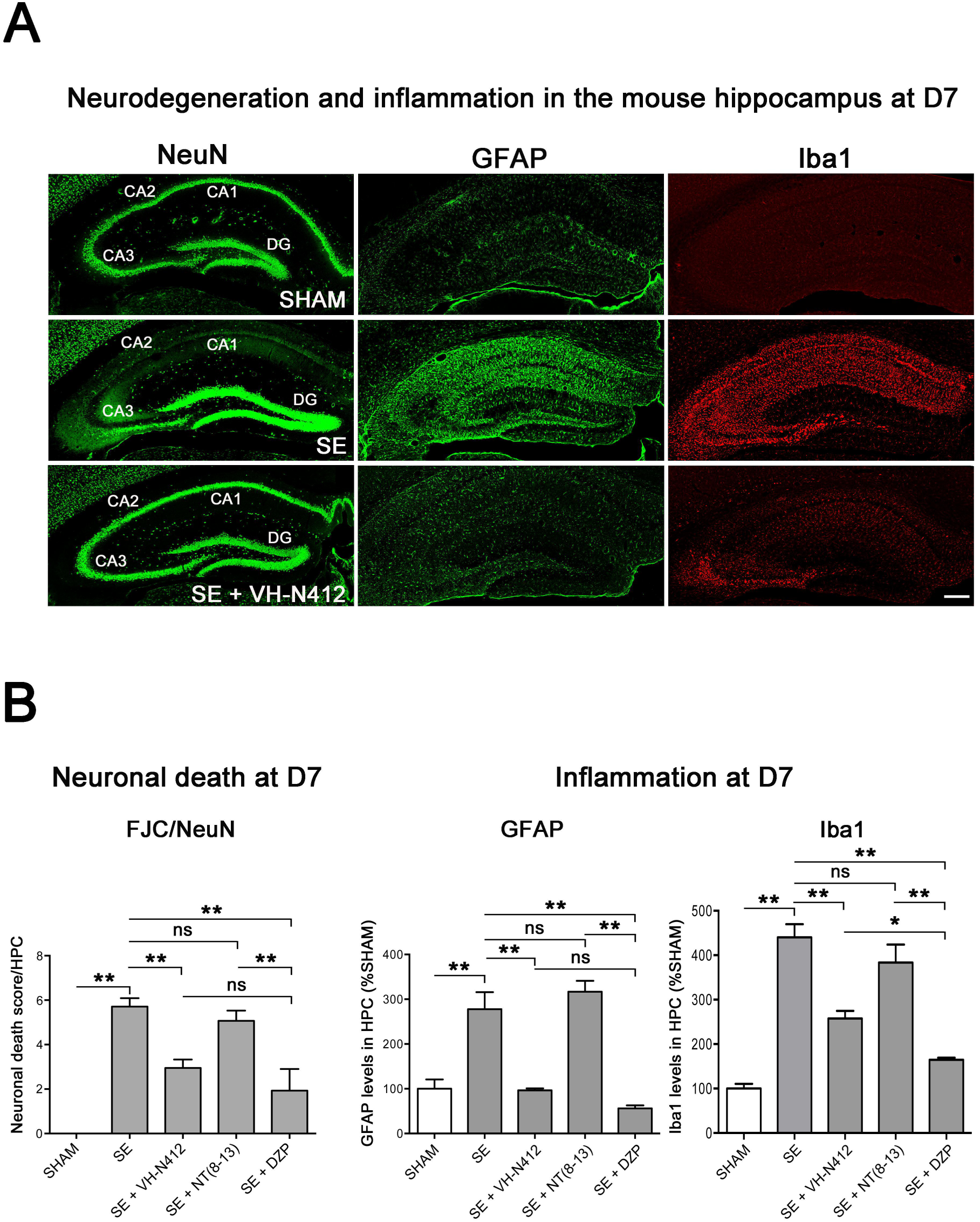
Neuroprotective and anti-inflammatory effects of VH-N412 following SE. A) Immunohistochemical labelling was used to assess the extent of brain damage in coronal sections of the dorsal hippocampus from SHAM, SE, SE + VH-N412, SE + NT (8-13) and SE + DZP animals at D7 post-SE. Left panels show neurons labelled with the anti-NeuN antibody directed against a neuronal specific nuclear protein in all animals. Middle and right panels show inflammation assessed with anti-GFAP and Iba1 antibodies to monitor astrocytic and microglial reactivity respectively. Scale bar: 200 μm in all panels. In SHAM animals, a basal labelling for GFAP and Iba1 was detected in the hippocampus. In SE animals, a strong activation of glial cells occurred in all hippocampal layers. This inflammatory response was nearly abolished when VH-N412 was administered 30 min after SE onset. B) Histograms comparing the mean neuronal death score, the mean GFAP and Iba 1 levels in the dorsal hippocampus of SHAM, SE, SE + VH-N412, SE + NT(8-13) and SE + DZP animals. NeuN and FJC labelling were used to quantify neuronal death and the effects of VH-N412 (left histogram). The neuronal death score was expressed as the mean scores ± SEM. GFAP and Iba1 labelling levels allowed quantification of glial inflammation, which was expressed as the average percentage ± SEM normalized to CTL SHAM. In SHAM animals, no neuronal death was observed in the hippocampus (score 0). In SE animals, significant neuronal death was observed in CA1-3 pyramidal cell layers and H. Neurodegeneration observed in SE animals, was significantly decreased when VH-N412 or DZP were administered 30 min after SE onset, but no changes were observed when NT(8-13) was administered. Asterisks indicate statistically significant differences: \**p*<0.05, ** *p*<0.01 (Tukey’s-test).

#### Effects of VH-N412 on glial-mediated inflammatory response

We used GFAP- and Iba1-immunolabelling to evaluate the effects of VH-N412 on astroglial-(Figure 6A, in green, middle panels) and microglial-(Figure 6A, in red, right panels) reactivity respectively. In SHAM animals, basal labelling for GFAP (100 ± 20.77%) and Iba1 (100 ± 18%) was visible in the hippocampal formation (Figure 6B, middle histogram). In SE animals, a significant activation of glial cells occurred in all hippocampal areas (GFAP: 277;74 ± 37.96%, 178%; *p*<0.01, Tukey’s test; Figure 6B, middle histogram; Iba1: 440.04 ± 29.86%, 340%; *p*<0.01, Tukey’s test; Figure 6B, right histogram). This inflammatory response was significantly decreased when VH-N412 (GFAP: 96.44 ± 4.48%; 181.50%; *p*<0.01, Tukey’s test; Figure 6B, middle histogram; Iba1: 257.69 ± 16.96%, 182.31%; *p*<0.01, Tukey’s test; Figure 6B, right histogram) or DZP (GFAP: 56.32 ± 6.61%; 221.41%; *p*<0.01, Tukey’s test; Figure 6B, middle histogram; Iba1: 164.59 ± 4.88%, 275.41%; *p*<0.01, Tukey’s test; Figure 6B, right histogram) were administered at SE30. No significant changes were observed when NT(8-13) was administered at SE30 (GFAP: 316.87 ± 24.36%; *p*>0.05; ANOVA; Figure 6B, middle histogram; Iba1: 383.32 ± 40.40%; *p*>0.05; ANOVA; Figure 6B, right histogram). Altogether, these results indicate that SE-induced neuroinflammation is partially prevented by VH-N412 treatment.

#### Effects of VH-N412 on mossy fiber sprouting

Temporal lobe epilepsy is associated with sprouting of the mossy fibers in the inner molecular layer (IML), in response to hilar cell loss (Jiao and Nadler, 2007; Sloviter et al., 2006). To further investigate the neuroprotective effect of VH-N412, we evaluated the extent of mossy fiber sprouting 8 weeks after SE. Since mossy fiber terminals are highly enriched in zinc ions, we used immunohistochemical labelling for the zinc vesicular transporter 3 (ZnT-3) to detect mossy fiber sprouting as illustrated in Figure 7. In all SHAM animals, mossy fiber terminals are present in the hilus-region and no terminals were observed in the GCL and IML of the DG (Figure 7A). In SE animals, mossy fiber terminals were not only observed in the hilar region as in SHAM mice, but also within the IML and GCL of the DG (Figure 7A). In comparison with SE animals (50.38 ± 15.57), the number of terminals innervating the IML was significantly reduced in animals administered with VH-N412 at SE30 (7.79 ± 2.54, 84.53%, *p*<0.01, Tukey’s test, Figure 7B) but was not significantly different when NT(8-13) (25.85 ± 6.39, *p*>0.05; ANOVA) was administered at SE30 (Figure 7B). Altogether, these results indicate that SE-induced mossy-fiber sprouting is partially prevented by VH-N412 treatment.

**Figure 7:**
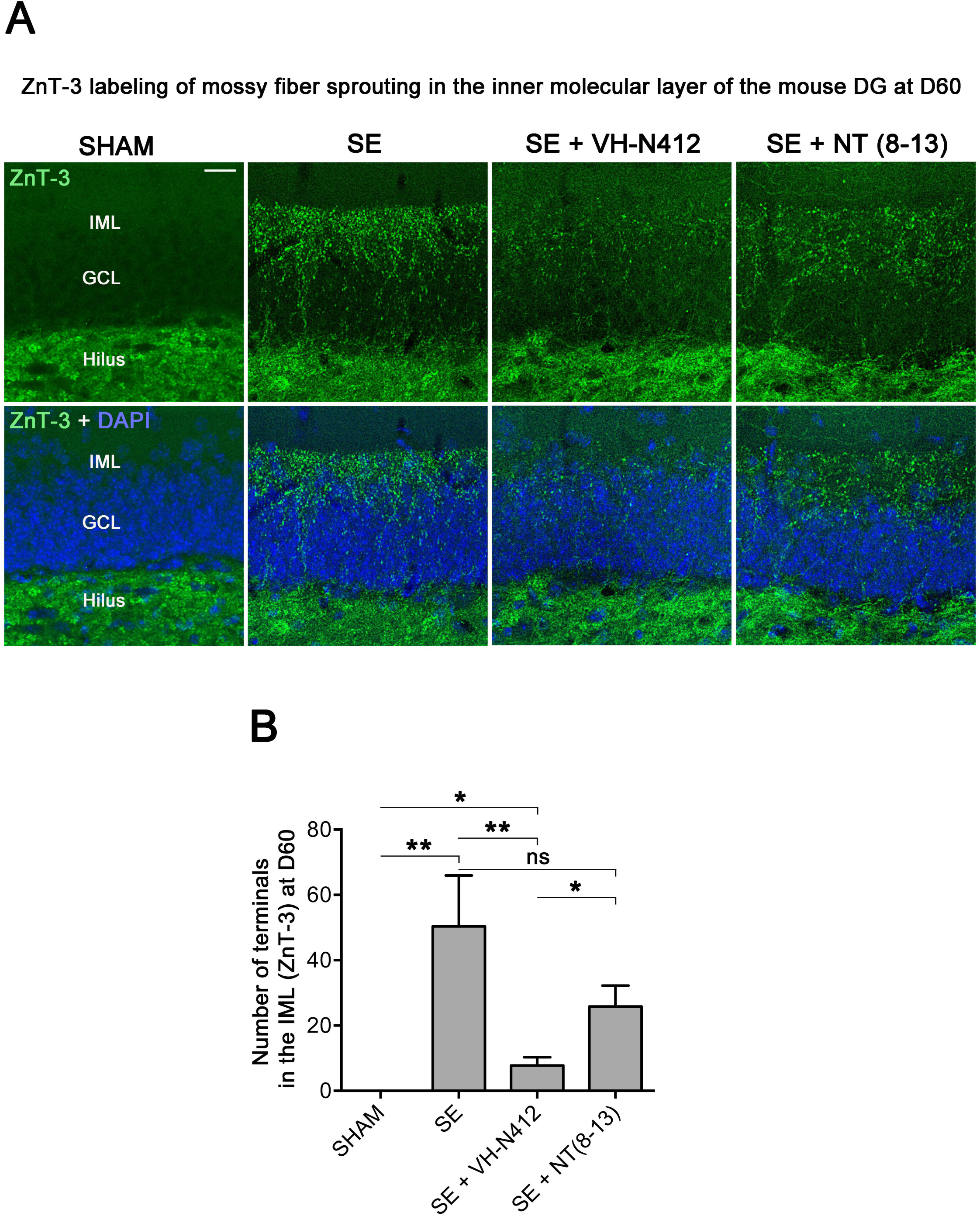
VH-N412 reduces mossy fiber sprouting in the hippocampus 8 weeks (D60) after SE. A) The effects of VH-N412 on mossy fiber sprouting was assessed 8 weeks after induction of SE with immunohistochemical labelling for the zinc vesicular transporter 3 (ZnT-3). In SHAM animals, mossy fiber terminals were only present in the H. In SE animals, in addition to ZnT-3 staining present in the H, mossy fiber terminals were also observed within the IML (A). Scale bar: 20 μm in all panels. B) Semi-quantitative analysis revealed that ZnT-3 staining in the IML at D60 was significantly reduced in animals administered with VH-N412 30 min after SE onset but unchanged when NT(8-13) was administered. Asterisks indicate statistically significant differences: \**p*<0.05, ** *p*<0.01 (Tukey’s-test).

### Effects of VH-N412 on learning and memory following SE

Epilepsy is associated with learning and memory difficulties in patients and animal models (Giovagnoli and Avanzini, 1999; Löscher and Stafstrom, 2023). We thus assessed whether VH-N412-induced reduction of neurodegeneration and inflammation ameliorates mnesic capacities of our epileptic animals. Three weeks after SE, a group of mice was submitted to behavioral tests (Figure 8). We first evaluated hippocampus-dependent learning and memory performance using the olfactory tubing maze, a test particularly well suited for the FVB/N mouse strain we used, with poor or deficient vision (Girard et al., 2016). Using the OTM test, the inter-trial interval (ITI) showed no significant effect across the five training sessions (MANOVA: F(8,92) = 1.16, non-significant (ns) and between the 3 groups (MANOVA: F(2,23) = 1.85; ns) (Figure 8A). However, considering the percentage of correct responses (MANOVA: F(8,92) = 0.99; ns), a subsequent group difference was observed between the 3 groups (MANOVA: F(2,23) = 6.23; *p*<0.01) (Figure 8B). Post-hoc analysis using the Newman-Keuls test revealed that SE mice reached a significantly lower percentage of correct responses in comparison with the two other groups (*p*<0.05). Selective ANOVAs showed a significant difference starting from the third training session between SE and SE + VH-N412 groups (ANOVAs: F(1,14) ≥ 5.12; *p*<0.05). In addition, a similar difference was also observed between SHAM and SE mice on session five (ANOVA: F(1,14) = 8.1; *p*<0.05) while no significant differences were observed between control and SE + VH-N412 mice (ANOVAs: F(1,18) ≤ 2.5; ns) during each of the 5 sessions.

**Figure 8:**
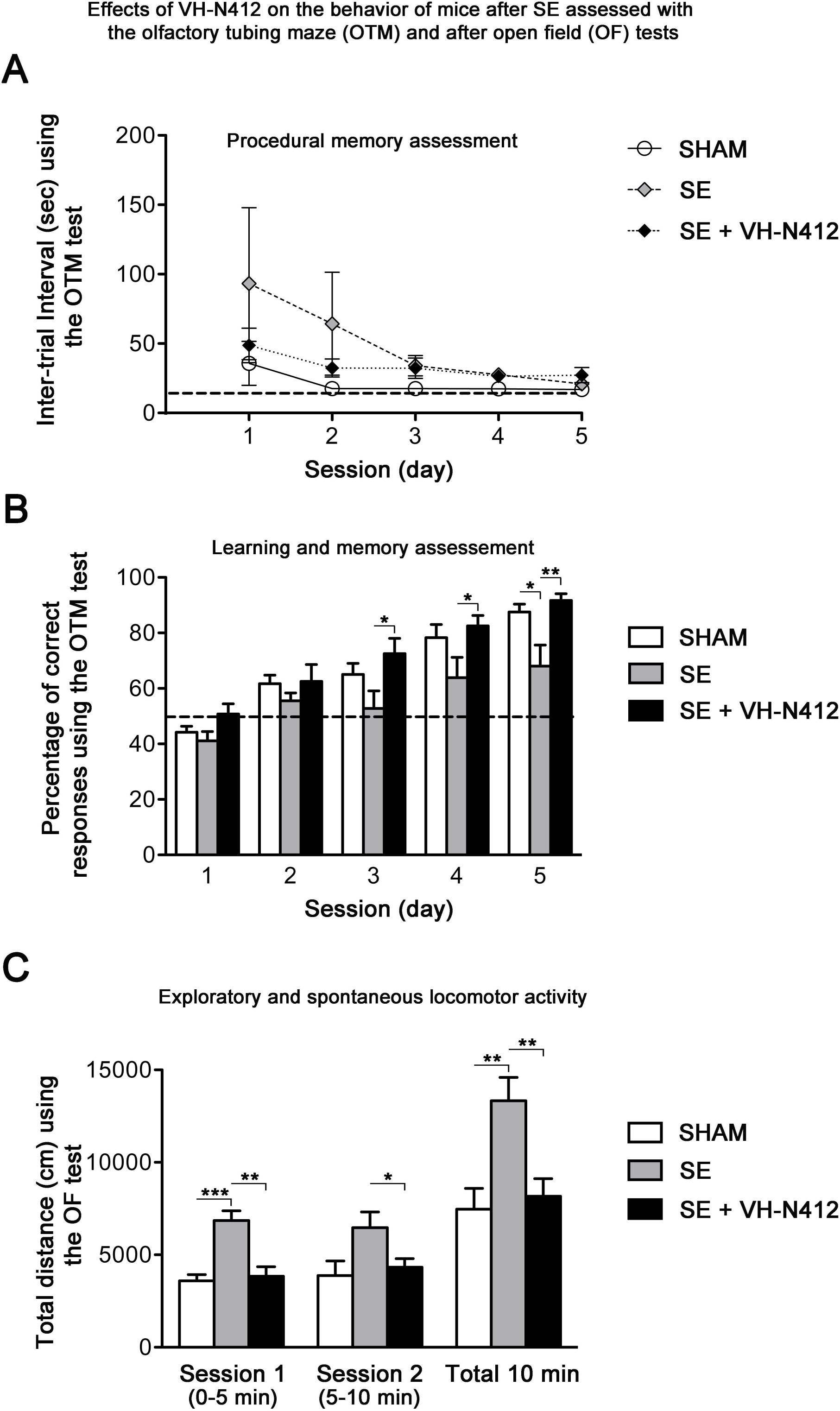
Effects of VH-N412 on the regulation of RBM3 and CIRBP mRNAs/proteins in the hippocampus and *in vitro*. A) Histograms related to analyses of RBM3 and CIRBP mRNA levels in the hippocampal tissue of CTL and mice treated with VH-N412 at 4, 8, and 16 H by RT-qPCR. Note that VH-N412 treatment increased the expression of RBM3 at 16 H while CIRBP peaked at 8 H compared to CTL mice. B) Western blots of CIRBP and RBM3 protein expression in VH-N412-treated mice at the same time points as above. Note that CIRBP and RBM3 are expressed predominantly at their expected molecular weight (MW) of ∼18 kDa (stars). However, bands of higher MW, between ∼22 kDa (RBM3) to ∼25 kDa (CIRBP) were also detected in agreement with (Rosenthal et al., 2017; Zhu et al., 2024). Similar patterns of expression of the higher MW bands compared to the ∼18 kDa band strongly suggest that they correspond to either different isoforms for each protein or result from post translational modifications. Protein levels were normalized with β-actin (∼ 42 kDa, star) in all samples, and samples from mice treated with VH-N412 at 4 (n=3; lanes 4-6), 8 (n=3; lanes 7-9), and 16 H (n=3; lanes 10-12) were normalized with CTL mice treated with PBS (CTL PBS; n=3; lanes 1-3). The graphs correspond to quantification of the western blots. RBM3 is upregulated at 4, 8, and 16 H in VH-N412-treated mice, whereas CIRBP is upregulated at 4 H after treatment. These results indicate that *in vivo*, VH-N412-induced hypothermia is associated with a regulation of cold stress proteins. C) *In vitro* quantification of mRNA levels of RBM3 and CIRBP analyzed by RT-qPCR after treatment with different concentrations of VH-N412 (0,1, 1 and 10 μM) of cultured hippocampal neurons. VH-N412 did not regulate RBM3 and CIRBP mRNA at all concentrations used in our cultured hippocampal neurons.

Exploratory and spontaneous locomotor activity of the same mice was assessed in the open-field paradigm (Figure 8C). SE mice exhibited a strong hyperactivity in comparison with SHAM mice. The average distances covered by mice from these two groups were significantly different on the first session of 5 min (ANOVA: F(1,14) = 29.56; *p*<0.001) and on the total of the two sessions (ANOVA: F(1,14) = 11.18; *p*<0.01). Treatment of SE mice with VH-N412 maintained a locomotor activity similar to SHAM mice on the 2 successive sessions (ANOVAs: F(1,18) ≤ 0.24; ns) and significantly lower than untreated SE mice on each of the 2 sessions of 5 min (ANOVAs: F(1,14) ≥ 5.92; *p*<0.05) (Figure 8C). No significant difference was observed between groups in the time spent in the center of the maze (data not shown). In all, we showed that VH-N412 treatment following SE preserves learning and memory capacities and normal locomotor activity in mice.

### Expression and regulation of cold shock RBM3 and CIRBP mRNA and proteins

Cold-shock proteins such as the RNA binding protein RBM3 and cold-inducible RNA binding protein (CIRBP) are involved in diverse physiological and pathological processes, including circadian rhythm, inflammation, neural plasticity, stem cell properties, and cancer development (reviewed in Zhu et al., 2016). In particular, cooling and hibernation in animals induces expression of RBM3 and CIRBP (Shiina and Shimizu, 2020) and boosting endogenous RBM3 levels through hypothermia is neuroprotective (Ávila-Gómez et al., 2020). We hence questioned whether the hypothermic and neuroprotective effects of VH-N412 are associated with RBM3 and CIRBP regulation in brain. A group of 9 mice was administered VH-N412 at the dose of 4 mg/kg eq. NT leading to transient hypothermia. Mice were divided into 3 groups that were sacrificed at 4, 8 and 16 H post VH-N412 administration. Brains were rapidly extracted, cut in two for mRNA and protein analysis. In each hemi-section, the hippocampus was isolated and snap frozen. RT-qPCR analysis normalized with GAPDH on pooled samples (3 animals per time point) showed significant increase of mRNA encoding RBM3 (2.86 ± 0.33, 3-fold) and CIRBP (1.53 ± 0.56, 1.5-fold) relative to PBS injected control (1.00 ± 0.00) at the 16 H and 8 H time points, respectively (Dunnett’s test, *p*<0.05) (Figure 9A). Next, using western blot and antibodies that detect RBM3, CIRBP and β-actin used as an internal control, we evaluated in hippocampal samples whether increased mRNA levels translate into increased protein levels. Results showed significant increase of RBM3 protein levels (4H: 270.88 ± 5.12%, 2.7-fold, Dunnett’s test, *p*<0.05; 8H: 465.74 ± 70.05%, 4.65-fold Dunnett’s test, p<0.001; 16H: and 251.97 ± 49.34%, 2.5-fold, Dunnett’s test, p<0.05) and CIRBP protein levels (4H: 235.15 ± 13.02%, 2.35-fold, Dunnett’s test, p<0.001) relative to PBS injected control (100.00 ± 0.00%) (Figure 9B). However, CIRBP protein levels were not significantly altered at the 8 H (157.24 ± 24.55%) and 16 H (109.79 ± 10.15%) time points relative to PBS injected control (100.00 ± 0.00, *p*>0.05; ANOVA) (Figure 9B).

**Figure 9:**
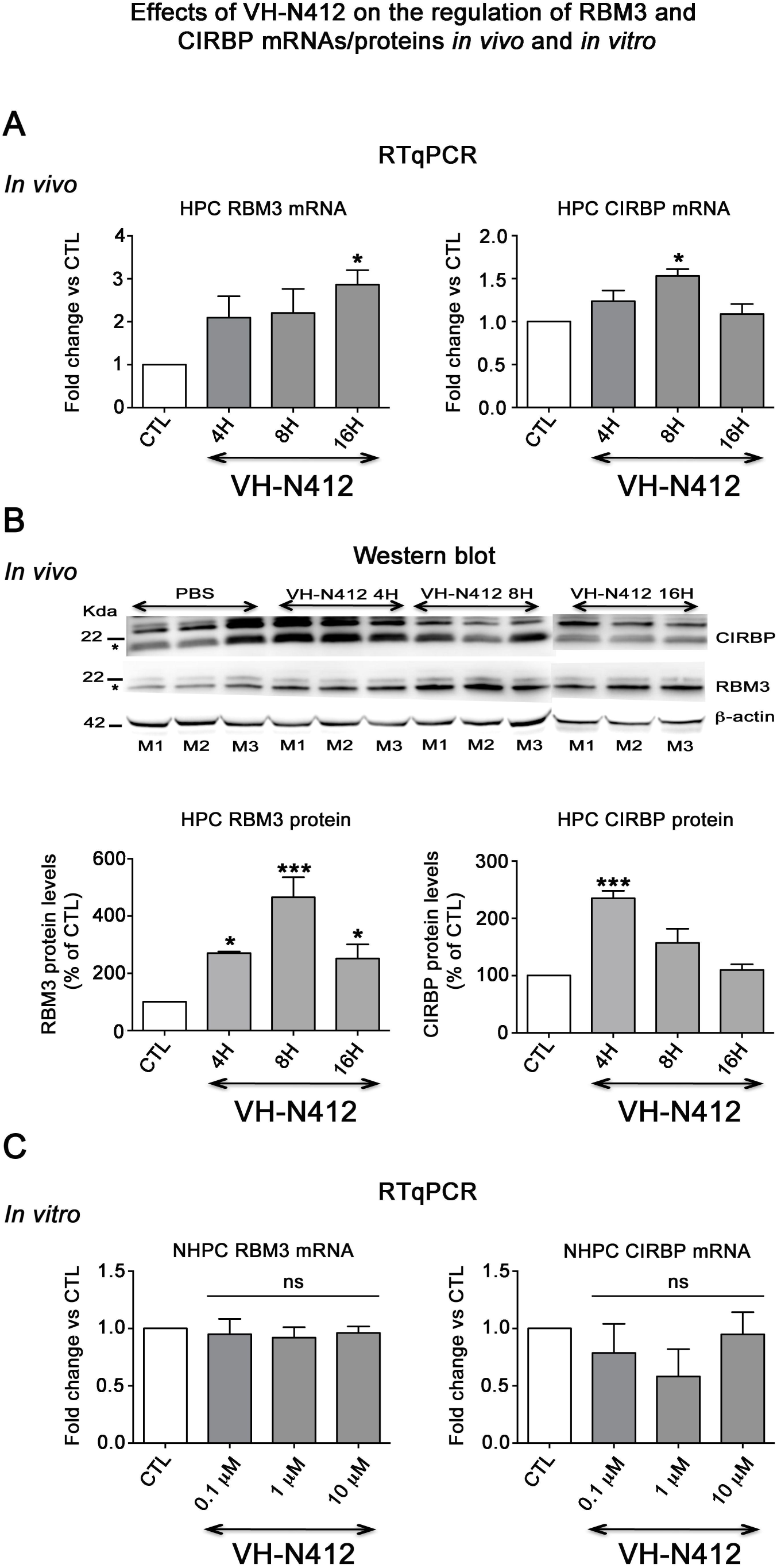
Following SE, VH-N412 restores hippocampus-dependent learning and memory and normal locomotor activity. A) and B) Learning and memory performance was assessed from 3-4 weeks after SE using the olfactory tubing maze (OTM) and C) locomotor activity using the open field (OF) test. A) illustrates the mean inter-trial interval (ITI) between the 12 trials in the OTM (in seconds, +/- SEM). The dashed line indicates the minimum fixed ITI (15 s). There was no difference between groups on the ITI. B) Mean percentage of correct responses obtained in the OTM during five training sessions of 12 trials per day. The dashed line denotes the chance level (%). From the second session, all animal groups had learned and memorized the test tasks. Only after the 5th session did the epileptic SE mice (n=6) show a significant impairment in memorization and learning while SE mice treated with VH-N412 showed similar performance to that of SHAM mice (n=10). C) illustrates the mean traveled distance in centimeters during 2 consecutive 5 min sessions (sessions 1 and 2) using OF test. SE mice (n=6) displayed a strong and significant hyperactivity in comparison with SHAM (n=10) while SE + VH-N412 mice (n=10) exhibited significant reduced exploratory and spontaneous locomotor activity, similar to that observed in SHAM mice.

To confirm that it is indeed hypothermia rather than VH-N412 that contributed to modulation of RBM3 and CIRBP mRNA levels *in vivo*, we incubated cultured primary rat hippocampal neurons with 3 concentrations of VH-N412 (0.1, 1 and 10 μM) for 24 H and assessed steady state levels of RBM3 and CIRBP mRNA by RT-qPCR. Our results showed that cultured neurons treated with VH-N412 did not display any significant differences of RBM3 and CIRBP mRNA levels at all concentrations compared to non-treated cultures (*p*>0.05; ANOVA, Figure 9C).

### Hippocampal expression of NTSR1 *in vitro* and *in vivo*

One important question raised by our results is whether NTSR1 receptors are indeed expressed in hippocampal neurons on which VH-N412 could exert some of its effects. To address this question, we performed immunocytochemistry and immunohistochemistry using a commercially available goat NTSR1 polyclonal antibody. We first validated the specificity of this antibody by using transfection experiments followed by immunocytochemistry in different cell types from different species, notably in human HEK 293 and 21-day *in vitro* (21 DIV) rat cultured hippocampal neurons (Figure 10).

**Figure 10:**
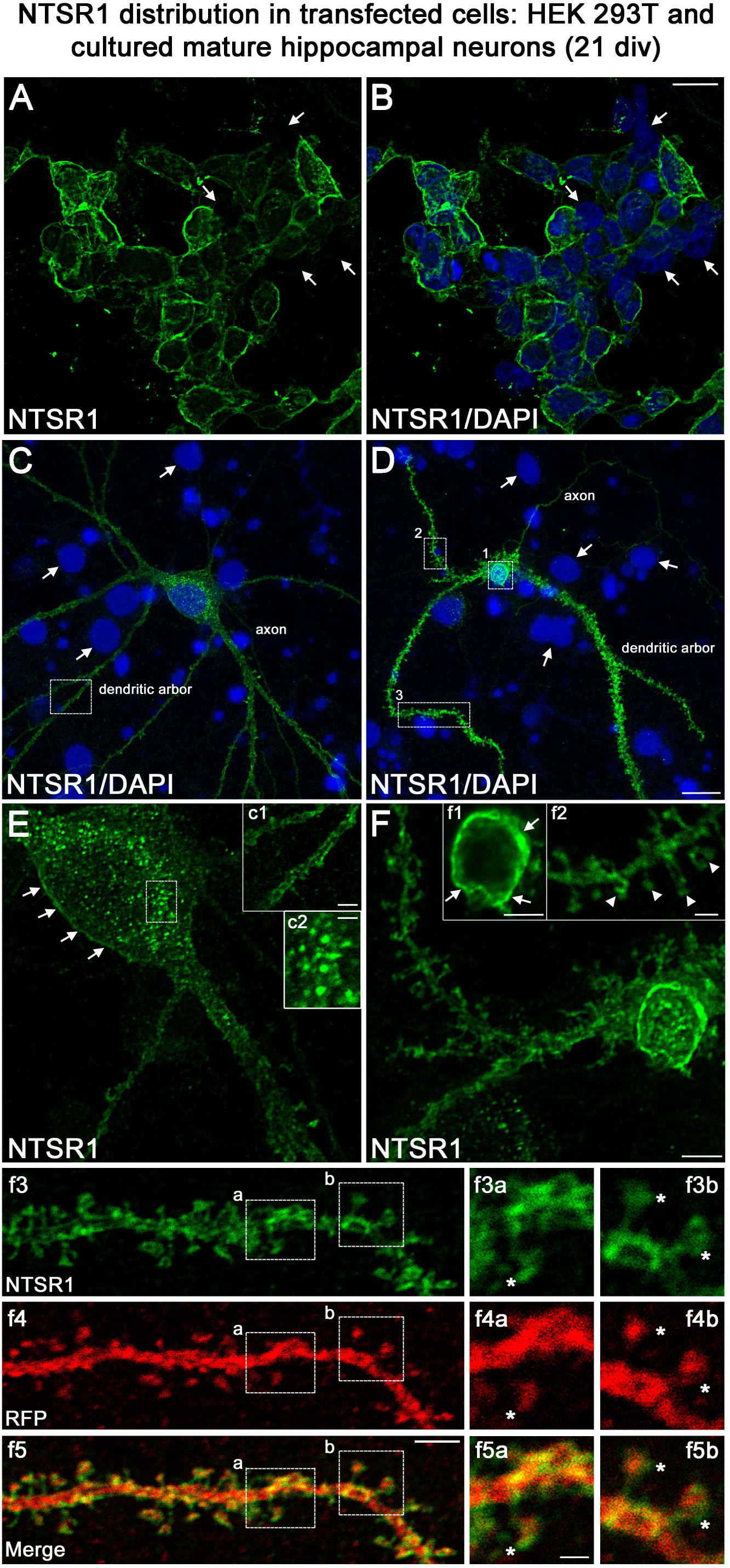
Validation of NTSR1 antibody. The specificity of the goat polyclonal NTSR1 antibody was assessed by using transfection experiments for 43 H followed by immunocytochemistry in different cell types including A and B) human HEK 293 cells and C and D) rat cultured mature hippocampal neurons (21 DIV). Cell nuclei were labelled with DAPI (blue). Both HEK 293 and hippocampal neurons displayed stronger NTSR1 immunolabelling (green) after transfection with a plasmid construct encoding rat NTSR1 (see cells double labelled for NTSR1 and DAPI), compared to non-transfected cells (see arrows, cells labelled for DAPI but not for NTSR1). Moreover, both types of cells exhibited high NTSR1 immunostaining within the cell body with a punctate pattern (see c2) and at the plasma membrane (see arrows in E, F, c1 and f1) as expected for receptor localization. The axons, the dendritic arbors, and their protuberances (see arrowheads in f2) of hippocampal neurons were also immunostained. f3 and f4 correspond to the dendritic portion of a neuron overexpressing NTSR1 (green) and RFP (red). RFP was used to outline the morphology of neurons including the dendrites and their dendritic spines. f5 corresponds to the merge of panels f3 and f4. Panels f3a and f3b correspond to the high magnification of NTSR1 labelling in 2 distinct areas of a dendrite (boxed in f3, f4, f5). Panels f4a and f4b correspond to RFP labelling in these same areas. Panel f5a corresponds to the merge of f3a and f4a. Panel f5b corresponds to the merge of f3b and f4b. Double immunostaining of NTSR1/RFP confirms that NTSR1 is located in dendritic spines. However, some of the NTSR1 immunolabelling is slightly shifted relative to RFP (see stars in panel f3a to f5b) suggesting NTSR1 localization in the cell membrane. Scale bars: 20 μm in A and B; 5 μm in C, D, E, F, c1, f1-f5. 2 μm in f3a-f5b.

Both HEK 293 cells (Figure 10A,B) and hippocampal neurons (Figure 10C,D) displayed higher NTSR1 immunolabelling (green) after transfection with a plasmid construct encoding rat NTSR1 compared to non-transfected cells (see arrows, cells labelled for DAPI but not for NTSR1). In addition, both types of cells exhibited high NTSR1 immunostaining within the cell body with a punctate pattern (c2 high magnification inset) and at the plasma membrane (arrows in E, F, c1, f1 high magnification insets), as expected for receptor localization. The axons, the dendritic arbors and protuberances of hippocampal neurons were also immunostained (Figure 10C,D, arrowheads in f2 high magnification inset), in agreement with (Boudin et al., 1998; Pickel et al., 2001). In the boxed areas, NTSR1 labelling (f3a,3b high magnification insets) and RFP (red), used to underline dendritic structures (f4a,4b high magnification insets), confirms that NTSR1 is also located in dendritic spines since NTSR1 and RFP proteins colocalized (f5a,5b high magnification insets). However, some of the NTSR1 labelling is slightly shifted relative to RFP, suggesting its localization at the cell membrane (stars in f3a-f5b high magnification insets).

#### Expression of endogenous NTSR1 in cultured hippocampal neurons

To assess the expression of endogenous NTSR1 and its localization, 21 DIV cultured hippocampal neurons were fixed and immunostained sequentially with the goat anti-NTSR1 polyclonal (red), the mouse anti-MAP2 antibody (blue) and a rabbit anti-drebrin E/A antibody (green) (Figure 11). Mature cultured hippocampal neurons displayed high levels of endogenous NTSR1 with a punctate pattern (Figure 11B and K, red) at high magnification of the boxed area in Figure 11A,F. Enlargement of the boxed area in Figure 11J and L illustrate that NTSR1 is closely apposed to dendritic shafts and dendritic spines, presumably at the level of the cell membrane, as revealed by the neuronal and dendritic shaft marker MAP2 (Figure 11E,H, blue), and dendritic spine marker drebrin (Figure 11I,H,J,L, green, arrows in high magnification insets). Note that no NTSR1 immunostaining was observed in filopodia (small arrow in Figure 11H,J,L).

**Figure 11:**
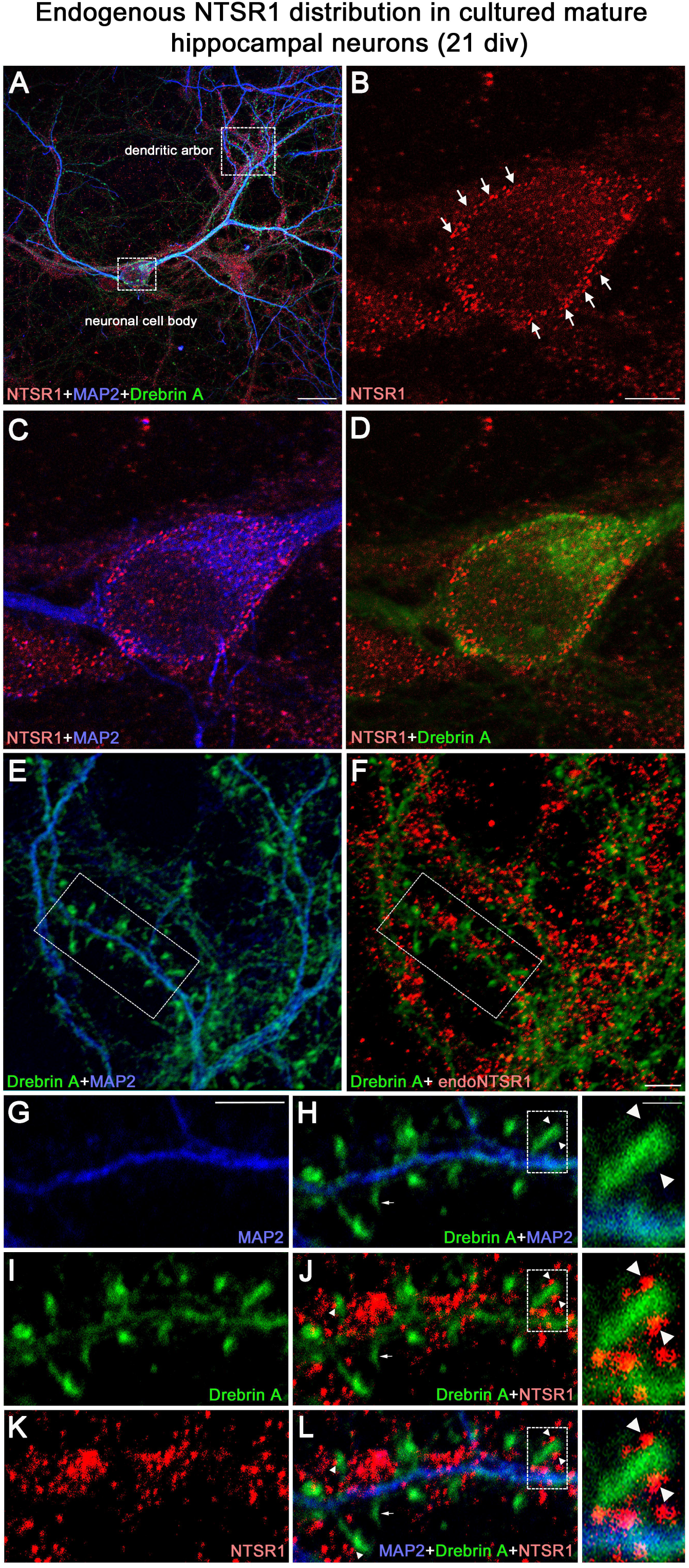
Expression of endogenous NTSR1 and its localization in mature cultures of hippocampal neurons. Twenty-one-day-old cultured hippocampal neurons were fixed and immunostained sequentially with antibodies against NTSR1 (red), MAP2 (blue) and drebrin E/A (green). Panels A, L and high magnification of the boxed area in L correspond to the merge of NTSR1/MAP2/drebrin. Panels E, H and F, J correspond to the merge of MAP2/drebrin and NTSR1/drebrin respectively. At high magnification of the boxed-in area in A (pyramidal neuron cell body) and F (dendrites), mature cultured hippocampal neurons displayed endogenous NTSR1 with a punctate pattern (B, C, D, F, J, K, L, red) similar to observations in transfected cells (Figure 10). Enlargement of the boxed-in area in F and J illustrate that NTSR1 (red) is closely apposed to dendritic shafts and dendritic spines, presumably at the level of the cell membrane, as revealed by neuronal and dendritic shaft marker MAP2 (E and H, blue), and dendritic spine marker drebrin (I, J and L, green, see arrows in high magnification insets). Note that no NTSR1 immunostaining was observed in filopodia (see small white arrows in H, J and L). Scale bars: 20 μm in A, 5 μm in B-L and 1 μm in boxed-in area in H, J and L panels.

#### NTSR1 is expressed in vivo, in adult mouse hippocampus

To confirm that the NTSR1 protein is expressed in the hippocampus *in vivo*, immunohistochemical labelling for NTSR1 was performed on control coronal sections of adult mice (3-5 months, Figure 12). At low magnification, NTSR1 immunolabelling was homogeneous in all areas and layers of the hippocampus (Figure 12A, DAPI in blue) and displayed a regional- and laminar-specific pattern within the hippocampus (Figure 12B, green). Moderate to strong NTSR1 immunolabelling was observed in all layers, including the stratum oriens (O), stratum radiatum (R) and the stratum lacunosum-moleculare (LM) of the CA1-CA2-CA3 areas, the stratum lucidum (SL) of CA2-CA3, the molecular layer (M) and H of the DG. The hippocampal pyramidal neurons (P) of CA1, CA2 and CA3 were relatively more immunostained as compared to GCL of the DG (Figure 12B). At high magnification, NTSR1 immunolabelling displayed a punctate pattern in CA1 (Figure 12C). NTSR1 was expressed at the plasma membrane of CA1 cell bodies (arrowheads in Figure 12D,F), consistent with the cell membrane localization of NTSR1 as well as within proximal dendrites of pyramidal neurons (arrows in Figure 12D,F). Note that similar to transfection results and immunocytochemistry on mature cultured neurons, the dendritic protuberances, reminiscent of dendritic spines, were also highly immunolabelled for NTSR1 (arrowheads in Figure 12D high magnification inset).

**Figure 12:**
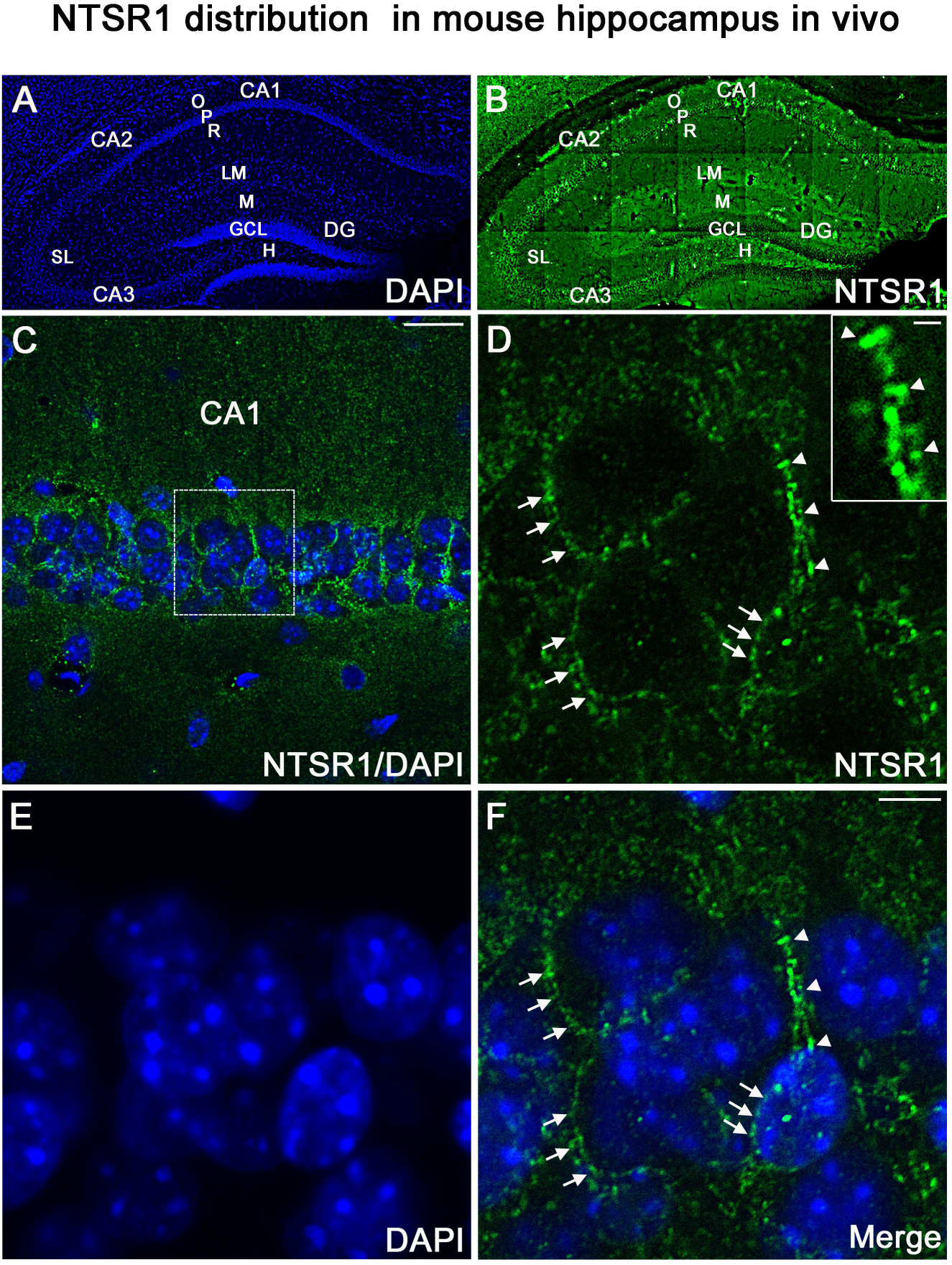
NTSR1 immunolabelling in mice hippocampal formation. A) and B) correspond to low magnification pictures showing coronal section of the mouse dorsal hippocampus processed with DAPI (blue), used to highlight the different cell layers of the hippocampus, and NTSR1 (green) antibody respectively. Moderate to strong NTSR1 immunolabelling levels were found in the stratum (O), (R), (LM), (SL), in pyramidal neurons of CA1, CA2 and CA3, M, H, and GCL of the DG. C) corresponds to the merge of DAPI and NTSR1 (green) of the CA1 area at high magnification. D-F) show high magnification of the boxed-in CA1 area illustrating pyramidal neurons immunolabelled with NTSR1 antibody (D, green) and counterstained with DAPI (E and F, blue). NTSR1 immunoreactivity is observed in the cell bodies, at the cell membranes (see arrowheads in E and F) as well as at the proximal dendrites of CA1 pyramidal neurons (see arrows in D and F). NTSR1 immunolabelling displayed a punctate pattern. Note that several dendritic protuberances displayed high levels of NTSR1 immunolabelling (see arrows in inset in D). Panel F corresponds to the merge of NTSR1/DAPI. Scale bars: 225 μm in A and B; 20 μm in C; 5 μm in D-F; 1 μm in inset in D.

### VH-N412 does not modulate neuronal hyperactivity induced by KA in hippocampal slices

The effects of VH-412 on seizure activity led us to question whether the conjugate could modulate hippocampal neuronal hyperactivity induced by KA. We addressed this question using acute hippocampal slices that were continuously perfused with ACSF pre-heated at 37°C. KA (300 nM) rapidly increased in the CA1 region the spontaneous firing rate that remained rather steady over the 110 min of KA exposure. The normalized firing rate was 0.86 ± 0.17 at the end of the experiment. When VH-N412 was applied at increasing doses of 0.1, 1 and 10 μM over a 30 min period, the KA-induced increase in firing rate in CA1 did not change significantly. Thus, the normalized firing rate was 0.96 ± 0.11 after 20-min exposure to 0.1 μM VH-N412, 0.92 ± 0.11 after 20-min exposure to 1 μM VH-N412 and 0.82 ± 0.13 after 20-min exposure to 10 μM VH-N412. A slight and transient increase of the firing rate was observed just after exposure to 0.1 μM VH-N412 for 2 out of the 4 recorded slices (Figure S2).

### Effects of VH-N412 on hippocampal neuronal survival and on dendrite length following NMDA or KA intoxication

Since SE-induced neurodegeneration was significantly decreased by VH-N412 (Figure 6A,B), we investigated *in vitro* whether these effects were associated with intrinsic neuroprotective effects of the conjugate. For this purpose, we induced intoxication of cultured hippocampal neurons by NMDA or KA treatment and evaluated whether VH-N412 elicited neuroprotection by assessing hippocampal neuronal survival (left histogram) and total dendrite length (right histogram). Note that the NTSR antagonists and VH-N412 alone showed no significant toxic effects on hippocampal neuronal survival and total dendrite length at all concentrations tested (Figure 13A). NMDA (100 µM, 10 min) induced significant neuronal death (51.77% of the CTL, *p*<0.001, Dunnett’s test, Figure 13B). VH-N412 at 1 µM and 10 µM protects significantly hippocampal neurons from NMDA-induced cell death compared to control (75.66% and 70.28% of the CTL, respectively, *p*<0.05, Dunnett’s test). These effects of VH-N412 were as potent as those of BDNF (50 ng/mL) used as a well-known neuroprotective molecule following NMDA intoxication (72.28% of the CTL, *p*<0.05, Dunnett’s test). In contrast to BDNF, we did not observe significant effect of VH-N412 on total dendrite length following NMDA intoxication (data not shown). Antagonizing NTSR by the SR142948A (0.1; 1 and 10 μM) and SR48692 (0.1 and 1 μM) NTSR antagonists blocked the neuroprotective effects mediated by VH-N412 in rat primary neuronal cultures injured by NMDA (Figure 13B).

**Figure 13:**
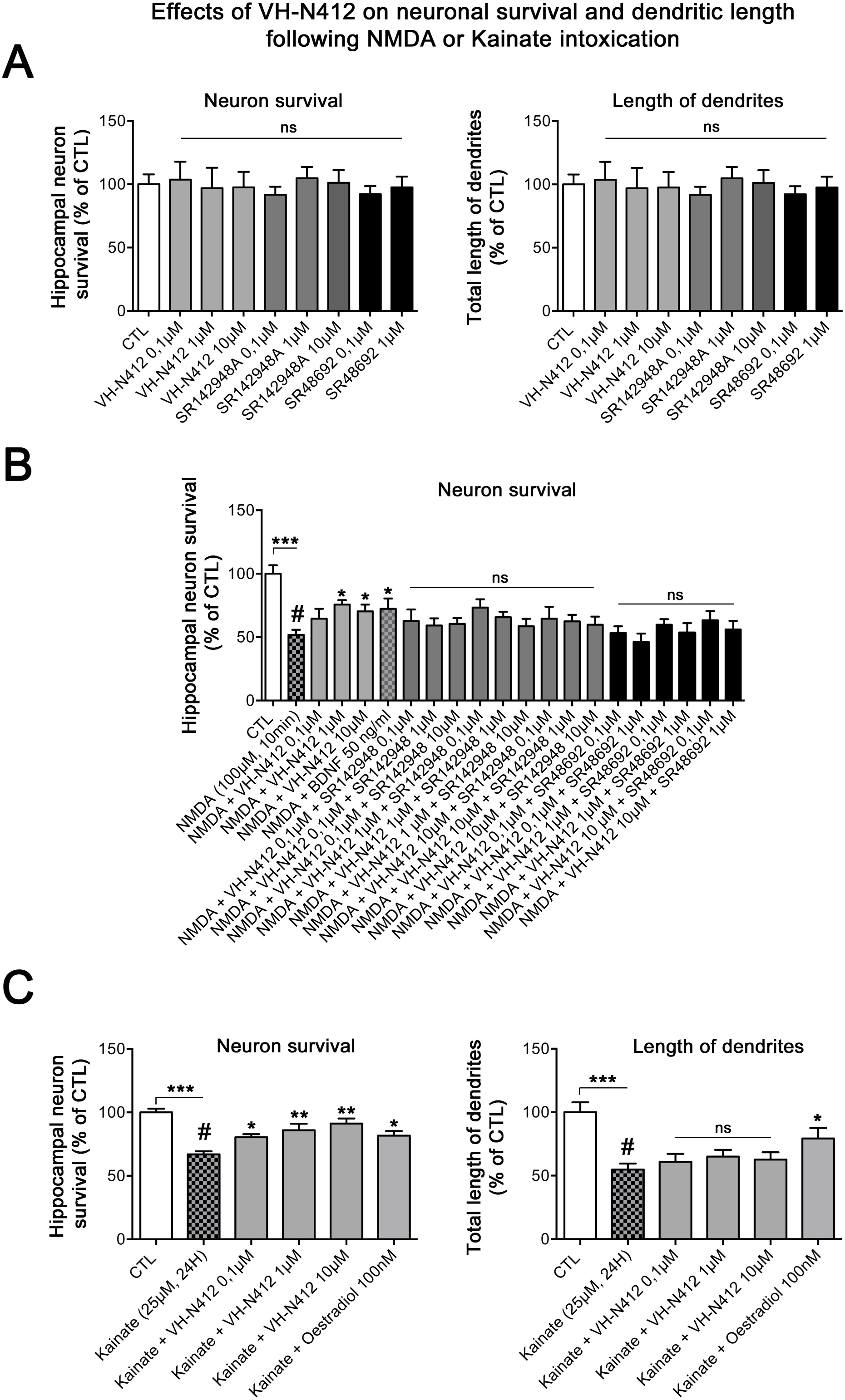
*In vitro* effects of VH-N412 on hippocampal neuronal survival and total dendrite length following NMDA or KA intoxication. A) Histograms illustrate the effects of VH-N412 and two NTSR antagonists, SR142948A and SR48692 on hippocampal neuronal survival (left histogram) and on total dendrite length (right histogram). These compounds alone showed no toxic effects on hippocampal neuronal survival and on total dendrite length at all concentrations used. B) illustrates the effects of VH-N412 on survival of primary hippocampal neurons injured by NMDA. VH-N412 promotes neuronal survival at 1 and 10 μM with similar potency as that of BDNF 50 ng/mL. Antagonizing NTSR by SR142948A and SR48692 blocks the neuroprotective effect of VH-N412 in the neuronal cultures. C) illustrates the effects of VH-N412 on hippocampal neuronal survival (left panel) and on total dendrite length (right panel) following KA intoxication (25 μM). VH-N412 promotes neuronal survival at all concentrations (0,1, 1 and 10 μM) with similar potency as that of Oestradiol at 100 nM. Following KA injury, VH-N412 did not display any significant effects on the total length of dendrites while Oestradiol did.

We next analyzed the effects of VH-N412 on hippocampal neuronal survival (Figure 13C, left histogram) and on total dendrite length (Figure 13C, right panel) following another neuronal intoxication model based on KA. As observed in Figure 13C, KA (25 µM, 24 H) induced significant neuronal death (66.88% of the CTL, p<0.001, Dunnett’s test). VH-N412 at 0.1, 1 and 10 µM significantly protected hippocampal neurons from KA-induced cell death compared to control (80.46%, *p*<0.05, 85.88%, *p*<0.01, 91.23%, *p*<0.01 of the CTL, respectively, Dunnett’s test). These effects of VH-N412 were similar to those of Oestradiol (100 nM) used as a well-known neuroprotective molecule following KA intoxication (81.68% of the CTL, *p*<0.05, Dunnett’s test). In contrast to Oestradiol, VH-N412 had no significant effects on the total length of dendrites following KA injury. Altogether, our data suggest that VH-N412 elicits neuroprotective effects by modulating hypothermia, as observed *in vivo*, but also displays intrinsic neuroprotective properties that are temperature independent.

## MATERIALS and METHODS

### Animals

All animals were housed six per cage in a temperature and humidity-controlled room (22 ± 2°C, 12 H light-dark cycles), had free access to food and water. Procedures involving animals were carried out according to National and European regulations (EU Directive N°2010/63) and to authorizations delivered to our animal facility (N° C13 055 08) and to the project (N° 00757.02) by the French Ministry of Research and Local Ethics Committee. All efforts were made to minimize animal suffering and reduce the number of animals used. Hypothermia was evaluated in 5 weeks old male Swiss CD-1 mice (Janvier laboratories, Le Genest-Saint-Isle, France) using a digital thermometer rectal probe (Physitemp Intruments Inc., Clifton, NJ, USA). Peptide stability was assessed in fresh mouse blood collected from Swiss CD-1 mice. For the Kainate (KA) model and for behavioral tests (see below) young adult male FVB/N mice (Janvier laboratories) were used. The FVB/N mice are reliable and well described mouse models of epilepsy, where seizures are associated with cell death and inflammation (Schauwecker, 2003; Wu et al., 2021).

### Peptide synthesis and characterization

All commercially available reagents and solvents were used as received without further purification. Resins (Fmoc-Gly-Wang Resin (100-200 mesh, 1% DVB, loading 0.7 mmol/g) and Fmoc-Leu-Wang Resin (100-200 mesh, 1% DVB, loading 0.7 mmol/g) were purchased from Iris Biotech (Marktredwitz, Germany), Bachem AG (Bubendorf, Switzerland), Sigma-Aldrich (St. Louis, MO, USA), Analytic Lab (St-Mathieu de Tréviers, France), Polypeptide Laboratories (Strasbourg, France) or ThermoFisher Scientific (Waltham, MA, USA). MS (Mass spectrometry) data were registered using a MALDI-TOF mass spectrometer (MALDI-TOF-TOF Ultraflex II Bruker, Mundelein, IL, USA) in positive mode. Internal calibration was applied by using 4-HCCA matrix m/z.

Reaction progress and purity monitoring were carried out using analytical RP-HPLC separation with a Dionex UltiMate^®^ 3000 system (ThermoFisher Scientific) equipped with a C18 Kinetex^TM^ (5 µm, 150 mm x 4.6 mm). Detection was performed at 214 nm. Elution system was composed of H_2_O/0.1%TFA (solution A) and MeCN/0.1%TFA (solution B). Flow rate was 2 mL/min with a gradient of 0-100% B in 4 min for reaction monitoring and 0-100% B in 10 min for purity assessment. Retention times (R_t_) from analytical RP-HPLC are reported in min. Crude products were purified by RP-HPLC with a Dionex UltiMate^®^ 3000 system equipped with a C18 Luna^TM^ (5 µm, 100 mm x 21.2 mm). Detection was performed at 214 nm. Elution buffer was composed of H_2_O/0.1%TFA (solution A) and MeCN/0.1%TFA (solution B). Flow rate was 20 mL/min.

### Synthesis of VH-N21 and VH-N41

#### Synthesis of Ac-[cMPRLRGC]c-G-OH

Peptide Ac-cMPRLRGC-G-OH (VH0445) was synthesized by solid phase peptide synthesis (SPPS) method on a CEM Liberty™ microwave synthesizer (Matthews, NC, USA), using standard Fmoc/tBu strategy and a Fmoc-Gly-Wang Resin (100-200 mesh, 1% DVB, loading 0.7 mmol/g) purchased from Iris Biotech on a 0.25 mmol scale. Fmoc deprotection was performed *via* micro-wave irradiation with piperidin 20% in DMF. Amino acids were coupled *via* micro-wave activation using the mixture aa/DIEA/HATU: 4/4/8 equivalent with respect to the resin. Coupling time was adjusted to 10 min. Double couplings were necessary for methionine and cysteine incorporation after the proline residue. N-terminal acetylation was performed manually using 50% acetic anhydride in DCM, 2 x 5 min at room temperature (RT) with gentle stirring. Resin-bound peptide was then cleaved using a solution comprised of TFA/TIS/H2O/EDT: 94/2/2/2 at RT for 2 H. A minimum of 15 mL of cleavage solution was used per gram of resin. Crude peptide was then precipitated using ice-cold ether, centrifuged at 3000 rpm for 8 min and lyophilized in H_2_O/0.1%TFA to obtain a white powder. Crude Ac-cMPRLRGC-G-OH peptide was then dissolved in AcOH 0.5% for a final concentration of 0.5 mg/mL. Ammonium carbonate (2N) was added to the peptide solution to reach an approximate basic pH of 8-9. K_3_[Fe(CN)_6_] (0.01N) was then added to the reaction mixture until a bright and persistent yellow color was observed. Monitoring of the reaction was performed by analytical RP-HPLC. After 30 min, the reaction mixture was purified by preparative RP-HPLC, R_t_) = 6 min, C18 Luna^TM^, 5 µm, 100 mm x 21.2 mm, flow rate = 20mL/min, phase A was 0.1% TFA in water, phase B was 0.1% TFA in acetonitrile, gradient was linear from 14 to 24% B over 30 min) to generate Ac-[cMPRLRGC]_c_-G-OH as a pure white powder (35.3 mg, 14 % yield, purity > 95%). R_t_= 1.73 min, C18 Kinetex^TM^, 5 µm, 150 mm x 4.6 mm, flow rate=2 mL/min, phase A was 0.1% TFA in water, phase B was 0.1%TFA in acetonitrile, gradient was linear from 0 to 100% B over 4 min). MALDI-TOF (*m/z*) for C_40_H_69_N_15_O_11_S_3_, [M+H]^+^ calc.1032.40 Da, found 1032.42 Da.

#### Synthesis of Ac-[cMPRLRGC]_c_-G-CH_2_-CH_2_-SH

Trt-cysteamine (2 eq.), PyBop (1.1 eq) and DIEA (4 eq.) were added to a 0.05 M solution of Ac-[cMPRLRGC]_c_-G-OH (28 mg) in anhydrous DMF. The reaction completion was monitored by RP-HPLC. The analysis showed the complete disappearance of the starting peptide after 5 min. DMF was then evaporated under vacuum to yield protected Ac-[cMPRLRGC]_c_-G-CH_2_-CH_2_-STrt as a pale-yellow oil. Trityl protection of the sulfur was removed by dissolving the crude oil in a solution of DCM/TIS/TFA: 3/1/1 (0.5 mL per mg of peptide). Reaction mixture was allowed to stir at RT. Monitoring of the reaction was performed by analytical RP-HPLC. DCM and TFA were then evaporated using inert gas bubbling and the crude peptide was subsequently precipitated with excess volume of ice-cold ether. After centrifugation at 3000 rpm for 8 min crude peptides were lyophilized in H_2_O/0.1% TFA and purified by preparative RP-HPLC (Rt = 16 min, C18 Luna^TM^, 5 µm, 100 mm x 21.2 mm, flow rate = 20 mL/min, phase A was 0.1% TFA in water, phase B was 0.1% TFA in acetonitrile, gradient was linear from 12 to 22% B over 30 min) to generate Ac-[cMPRLRGC]_c_-G-CH_2_-CH_2_-SH as a pure white powder (16 mg, 54% yield, purity > 95%). R_t_= 1.81 min (C18 Kinetex^TM^, 5 µm, 150 mm x 4.6 mm, flow rate=2 mL/min, phase A was 0.1% TFA in water, phase B was 0.1%TFA in acetonitrile, gradient was linear from 0 to 100% B over 4 min). MALDI-TOF (*m/z*) for C_42_H_74_N_16_O_10_S_4_, [M+H]^+^ calc. 1091.40 Da, found 1091.44 Da.

#### Synthesis of [Lys(MHA)6]NT

A solution of sulfo-EMCS (12.2 mg, 29.8 µmol, 1 eq.) in 2 mL of PBS 4X (pH=7.4) was added to a solution of commercial NT (pELYENKPRRPYIL-OH, 50 mg, 29.8 µmol, 1 eq.) in 7 mL PBS 4X (pH=7.6). Reaction mixture was allowed to stir at RT. Monitoring of the reaction was performed by analytical RP-HPLC. After overnight stirring the crude mixture was purified by preparative RP-HPLC (R_t_ = 15 min, C18 Luna^TM^, 5 µm, 100 mm x 21.2 mm, flow rate = 20mL/min, phase A was 0.1% TFA in water, phase B was 0.1% TFA in acetonitrile, gradient was linear from 21 to 31% B over 30 min) to generate [Lys(MHA)6]NT as a pure white powder (m=25 mg, 45% yield, purity > 95%). R_t_= 2.15 min (C18 Kinetex^TM^, 5 µm, 150 mm x 4.6 mm, flow rate=2 mL/min, phase A was 0.1% TFA in water, phase B was 0.1% TFA in acetonitrile, gradient was linear from 0 to 100% B over 4 min). MALDI-TOF (*m/z*) for C_88_H_133_N_22_O_23_, [M+H]^+^ calc. 1865.98 Da, found 1865.97 Da.

#### Conjugation of [Lys(MHA)6]NT to Ac-[cMPRLRGC]c-G-CH2-CH2-SH

[Lys(MHA)6]NT (1 eq) was added to a solution of Ac-[cMPRLRGC]_c_-G-CH_2_-CH_2_-SH in PBS 1X (1 eq., 0.003M, pH 7.4). The reaction mixture was allowed to stir at RT. Monitoring of the reaction was performed by analytical RP-HPLC. After reaction completion the mixture was purified by preparative RP-HPLC (R_t_ = 23 min, C18 Luna^TM^, 5 µm, 100 mm x 21.2 mm, flow rate = 20mL/min, phase A was 0.1% TFA in water, phase B was 0.1% TFA in acetonitrile, gradient was linear from 20 to 30% B over 40 min) to generate VH-N21 as a pure white powder (34% yield, purity > 95%). R_t_= 2.11 min (C18 Kinetex^TM^, 5 µm, 150 mm x 4.6 mm, flow rate=2 mL/min, phase A was 0.1% TFA in water, phase B was 0.1% TFA in acetonitrile, gradient was linear from 0 to 100% B over 4 min). MALDI-TOF (*m/z*) for C_130_H_206_N_38_O_33_S_4_, [M+H]^+^ calc. 2956.30, found 2956.67 Da. The molecule resulting from the conjugation of [Lys(MHA)6]NT to Ac-[cMPRLRGC]_c_-G-CH_2_-CH_2_-SH was named VH-N21. Another conjugate was also synthesized based on the conjugation of [Lys(MHA)6]NT to the previously described LDLR-binding peptide VH04129 [cM”Pip”RLR”Sar”C]_c_ (Jacquot et al., 2016), with the exception that the N-ter was propionylated rather than acetylated. This conjugate was named VH-N41.

### Synthesis of VH-N412

VH-N412 was synthesized by solid phase peptide synthesis (SPPS) method using standard Fmoc/tBu strategy and a Fmoc-Leu-Wang Resin (100-200 mesh, 1% DVB, loading 0.7 mmol/g) purchased from Iris Biotech on a 0.25 mmol scale. Synthesis was performed on a CEM Liberty™ microwave synthesizer except for the Fmoc-21-amino-4,7,10,13,16,19-hexaoxaheneicosanoic acid that was introduced manually overnight using COMU/DIEA (2/2/8 eq. with respect to the resin) in anhydrous DMF (0.2 M) as coupling cocktail. For automated synthesis Fmoc deprotection was performed *via* micro-wave irradiation with piperidin 20% in DMF. Amino-acids were coupled *via* micro-wave activation using the mixture aa/DIEA/HATU: 4/4/8 equivalent with respect to the resin. Coupling time was adjusted to 10 min. Double couplings were necessary for methionine and cysteine incorporation after the proline residue in the VH moiety. N-terminal propionylation was performed manually using 50% propionic anhydride in DCM (2 x 5 min at RT with gentle stirring). Resin-bound peptide was then cleaved using a solution comprised of TFA/TIS/H2O/EDT: 94/2/2/2 for 2 H at RT. A minimum of 15 mL of cleavage solution was used per gram of resin. Crude peptide was then precipitated using ice-cold ether, centrifuged at 3000 rpm for 8 min and lyophilized in H_2_O/0.1% TFA to obtain a white powder. Crude Pr-cMPRLRGC-PEG6-RRPYIL-OH peptide was then dissolved in AcOH 0.5% to reach 0.5 mg/mL final concentration. Ammonium carbonate (2N) was added to the peptide solution to reach an approximate basic pH of 8-9. K_3_[Fe(CN)_6_] (0.01N) was then added to the reaction mixture until a bright and persistent yellow color was observed. Monitoring of the reaction was performed by analytical RP-HPLC. After 30 min, the reaction mixture was purified by preparative RP-HPLC (Rt = 18 min, C18 Luna^TM^, 5 µm, 100 mm x 21.2 mm, flow rate = 20 mL/min, phase A was 0.1% TFA in water, phase B was 0.1% TFA in acetonitrile, gradient was linear from 19 to 29 % B over 30 min) to generate Pr-[cMPRLRGC]_c_-PEG6-RRPYIL-OH as a pure white to yellow powder (73.5 mg, 14% yield, purity > 95%). R_t_= 4.34 min (C18 Kinetex^TM^, 5 µm, 150 mm x 4.6 mm, flow rate=2 mL/min, phase A was 0.1% TFA in water, phase B was 0.1% TFA in acetonitrile, gradient was linear from 0 to 100% B over 10 min). MALDI-TOF (*m/z*) for C_94_H_163_N_27_O_24_S_3_, [M+H]^+^ calc. 2151.16, found 2151.11 Da.

### LDLR-binding using surface plasmon resonance analysis

Recombinant human LDLR (His-tagged) was purchased from Sino Biological (Beijing, China). Interaction of ligands with hLDLR was tested at 25°C using a Biacore T200 apparatus (GE Healthcare, Buc, France) and HBS (50 mM HEPES-NaOH pH7.4, 150 mM NaCl, 0.005% Tween-20, 50 µM EDTA) as running buffer. hLDLR was immobilized on a NiHC sensor chip (Xantec, Dusseldorf, Germany) at a density of around 25 pmol/mm2. Binding to hLDLR-coated flow cells was corrected for non-specific binding to uncoated flow cell. The single-cycle kinetic method was used to measure the affinity of the VH4129 peptide and vectorised NT (VH-N412) for hLDLR. VH4129 and VH-N412 were serially diluted 2-fold in running buffer yielding concentrations ranging from 50 to 800 nM and samples were injected sequentially every 2 min at 30 µl/min using increasing concentrations. Blank run injections of HBS were performed in the same conditions before injection of the ligands. Double-subtracted sensorgrams were globally fitted with the 1:1 binding model from Biacore T200 Evaluation version 2.0. Data are representative of at least 3 independent experiments.

### Binding to human and rat NTSR1 using a competition assay on NTSR1-expressing membrane extracts

The binding affinity of the neurotensin conjugates was assessed using a competition binding assay using rat and human NTSR1 membrane homogenates. The cell membranes enriched for the human NTSR1 were purchased from Perkin-Elmer (Villebon-sur-Yvette, France) and used according to manufacturer’s recommendations. The rat NTSR1 membranes were prepared as follows: one day before transfection, HEK 293 cells (ECACC, Salisbury, UK) were seeded and cells were transfected using Lipofectamine 2000 (Life Technologies, Saint Aubin, France) with the rat NTSR1 plasmid construct (Origene, Rockville, Maryland, USA). Cell expression of the rat NTSR1 receptor was evaluated using western blot and immunocytochemistry as described in detail below. At 100% confluence, the medium was removed, and the cells were harvested using PBS EDTA 5 mM buffer. They were washed with ice-cold PBS and centrifuged at 1,700 g for 10 min at 4°C. The pellet was suspended in ice-cold buffer (1 mM EDTA, 25 mM sodium phosphate, 5 mM MgCl_2_, pH 7.4) and homogenized using a Potter-Elvehjem homogenizer (Fisher Scientific, Elancourt, France). The homogenate was centrifuged at 1,700 g for 15 min (4°C). The pellet was washed, resuspended, and centrifuged at 1,700 g for 15 min (4°C). The combined supernatants were centrifuged at 100,000 g for 40 min (4°C) on a TL110 rotor (Beckman Coulter, Paris, France) and the pellet was suspended in the same buffer. Protein concentrations were determined according to the adapted Lowry method (Biorad, Hercules, CA, USA). The membrane preparations were aliquoted and stored at −80°C. Assays on hNTSR-1 and rNTSR-1 were performed using membrane homogenates at a final concentration of 0.5 µg/well or 0.75 µg/well, respectively, and the radioligand [^3^H]-neurotensin (specific activity 99.8 Ci.mmol^-1^; Perkin-Elmer) at a concentration of 3 nM. Specific binding of the radioligand was determined with K_D_ values of 0.81 nM and a B_max_ of 34 pmol per mg membrane protein for binding to hNTSR-1 and K_D_ values of 0.93 nM and a B_max_ of 2.5 pmol per mg membrane protein for binding to hNTSR-1. Non-specific binding was determined in the presence of 5 µM NT. Each assay was performed in a 96-well plate in a total reaction volume of 100 μL in binding buffer (50 mM Tris-HCl pH 7.4, 0.1% bovine serum albumin (BSA, Sigma). The assay was subsequently incubated for 24 H at RT. The content of each well was rapidly filtered through Unifliter®-96, GF/C® filters (Perkin-Elmer) presoaked with 25 µL of 0.5 % polyethylenimine) using a MicroBeta FilterMate-96 Harvester (Perkin-Elmer) and each well was rinsed 5 times with washing buffer (10 mM Tris-HCl, pH 7.2). Radioactivity (cpm) of each dried filter was measured by adding 25 µL of MicroScint™-O (Perkin-Elmer) and quantified using TopCount NXT™ Microplate Scintillation and Luminescence Counter (Perkin-Elmer). Specific binding was typically 90% or greater of the total binding. Dose-response curves were plotted using KaleidaGraph to determine IC50 values. Assays were performed in duplicate. Ki values were determined from mean IC50s using the Cheng-Prusoff conversion.

### *In vitro* stability of conjugates in freshly collected blood

*In vitro* blood stability (half-life or t_1/2_) of the VH-N21 and VH-N412 conjugates was assessed and compared to that of NT alone as previously performed with free VH0445 analogues (Malcor et al., 2012; Jacquot et al., 2016). Briefly, each peptide was incubated at the nominal concentration of 2 µM up to 3 H at 37°C in freshly collected Lithium Heparin blood from Swiss CD-1 mouse or human. The analyte was quantified in the plasma fraction at several time-points by liquid chromatography-tandem mass spectrometry (LC-MS/MS), using a Shimadzu LC equipment coupled to an API 4000 triple quadrupole mass spectrometer (Applied Biosystems, Foster City, CA, USA). *In vitro* half-life (t_1/2_) was estimated from the logarithmic regression of each kinetic profiles (first-order reaction kinetics: C(t)=C_0_.e^-kt^) and given by t_1/2_=ln2/k.

### Radiolabelling of vectorised and non-vectorized NT and *in situ* brain perfusion in mice

Radiolabelled activated NT compounds were obtained either by coupling [3H]Tyr3-NT with the LDLR-binding peptide or by coupling of a tritiated LDLR-binding peptide to NT molecules. NT was radiolabeled by palladium-catalyzed dehalogenation of 3,5diiodoTyr3-NT using tritium gas and Pd/C while the LDLR-binding peptide was radiolabeled by coupling of a tritiated propionyl succinimide ester to its N-terminus. The specific radioactivity (SRA) was typically in the range of 50 Ci/mmol. The total quantity of radioactivity prepared for each synthesis was generally between 100 and 1000 µCi.

BBB transport of the NT and NT-peptide conjugates was assessed in adult male mice as described in detail in Dagenais et al. (2000). Briefly, the *in situ* brain perfusion of the radiolabelled compounds was performed during 120 s with a perfusate flow rate of 2.5 mL/min. The initial transport was expressed as the brain clearance K_in_ and corrected for the vascular volume with ^14^C-sucrose co-perfusion. ^14^C-sucrose, a compound that does not cross the intact BBB, allowed assessment of its physical integrity. The BBB transport of the radiolabeled peptides was expressed as the Kin (μL.s^-1^.g^-1^) = V_brain_/T (where V_brain_ is the apparent volume of distribution of the tritium labelled compound corrected from vascular contamination, and where T is the perfusion time).

### Kainate model of temporal lobe epilepsy

Forty-eight young adult male FVB/N mice (25-30 g; 9-10 weeks-old) were injected s.c. with a single dose of KA (45 mg/kg; Abcam, Cambridge, UK) to generate mice with spontaneous recurrent seizures as a hallmark of status epilepticus (SE), as previously described (Schauwecker and Steward, 1997; Gröticke et al., 2008; Wu et al., 2021). KA-injected mice were individually housed and received a 0,5 mL i.p. dose of glucose G5 and had free access to agarose-doliprane gel (2.4%; Sanofi, Gentilly, France) to avoid pain. Mice were observed during 9 H for onset and extent of seizure activity. Seizure activity was scored by visual inspection according to a modified Racine (1972) scale: stage 0: exploration, “normal behavior”; stage 1: immobility, staring; stage 2: head nodding and/or extended tail; stage 3: forelimb clonus and/or circling behavior; stage 4: rearing with forelimb clonus and falling; stage 5: continuous rearing and falling; stage 6: severe tonic-clonic seizures. Only animals having displayed at least stage 5 were included in the study and 5 animal groups were generated: 1) animals injected 30 min after the onset of the SE with one intravenous (tail vein) bolus injection of the VH-N412 compound at the dose of 4 mg/kg eq. NT (group “SE30 + VH-N412”, n=13); 2) animals injected 30 min after the onset of the SE with an intravenous bolus injection of NT at the dose of 4 mg/kg eq. NT (group “SE30 + NT(8-13)”, n=14); 3) animals injected 30 min after the onset of the SE with DZP i.p. (Valium®, Roche, Basel, Switzerland) at the dose of 15 mg/kg (group “SE30 + DZP”, n=5); 4) animals injected 30 min after the onset of the SE with saline 0.9% (vehicle control) (group “SE”, n=11); 5) a negative control group of age-matched mice was administered saline 0.9% alone (group “SHAM”, n=5). All animals were weighed and monitored daily. The measurement of body temperature was performed using a digital thermometer rectal probe before KA injection and every 30 min during 3 H thereafter. Seizure intensity score was assessed following KA injection, and every 30 min during 3 H. As SE did not occur at the same time for each animal, recordings were performed from the onset of SE for comparative analysis of changes in body temperature and for seizure intensity score.

To study mice in the chronic stage of epilepsy with spontaneous seizures, they were observed daily (at least 3 hours per day) for general behavior and occurrence of SRS. These are highly reproducible in the mouse KA model, allowing for visual monitoring and scoring of epileptic activity. After 3 weeks, most animals exhibited SRS, with 2 to 3 seizures per day, similar to previous observations (Wu et al., 2021). The detection of at least one spontaneous seizure per day was used as criterion indicating the animals had reached chronic phase that can ultimately be confirmed by mossy fiber sprouting.

### Tissue preparation

Mice were deeply anesthetized with pentobarbital (700 mg/kg; ProLabo, Fontenay-Sous-Bois, France) and transcardially perfused with 4 % paraformaldehyde. The brains were extracted and post-fixed for 1 H at RT and rinsed in 0.12M phosphate buffer (0.12M PB; pH 7.4). Forty μm coronal sections were cut on a vibratome, immersed in a cryoprotective solution and stored at −20°C until histological assessment. Every tenth section was stained with cresyl violet to determine the general histological characteristics of the tissue within the rostro-caudal extent of the brain. From each mouse, selected sections from the dorsal hippocampus were then processed for tissue evaluation by using Neuronal nuclear antigen (NeuN) immunohistochemistry or FJC staining. Adjacent sections were treated for assessment of inflammation and mossy fiber sprouting. Histopathological analyses were performed at two post-SE intervals: at 1 week to estimate hippocampal sclerosis and at 2 months to assess mossy fiber sprouting. For each experiment, sections from the different animal groups were processed in parallel.

### Fluoro-Jade C staining and scoring of neurodegeneration

To estimate cell degeneration, we performed FJC histo-staining (Chemicon, Temecula, CA, USA) on sections of dorsal hippocampus from SHAM, SE, SE + VH-N412, SE + NT(8-13) and SE + DZP animals according to the standard protocol of Schmued et al. (2005) and Bian et al. (2007). This histostaining does not depend on the mode of cell death (Ikenari et al., 2021). The specimens were analyzed using a confocal microscope and images were acquired using Zen software (Zeiss, LSM 700, Jena Germany), and processed using Adobe Photoshop and *ImageJ* softwares. The mean fluorescence intensity of FJC (mean grey value) was measured in Z-stack x10 z1 acquired images of pyramidal cell layers CA1, CA3, H, GCL. A minimum of 4 slices were analyzed on both sides of the dorsal hippocampus for each mouse of each group. Values were obtained following analysis of n=3-4 mice per group. We subtracted fluorescence value in areas devoid of stained cells in the same sections; an average value of 4 for each area was obtained from every image. Data are presented as the mean percentage of fluorescence of FJC normalized to that of SHAM (mean ± SEM).

To confirm the neuronal identity of degenerating cells and to evaluate the degree of neurodegeneration we scored NeuN and FJC staining on hippocampal sections. The score ranged from 0 (absence of neuronal death or FJC positive neurons) to 7 (maximal neurodegeneration). A score of 1 was given for positive neurons either in the H, in the GCL, in the CA3, CA1 or CA2 pyramidal cell layers. Whenever an intense neurodegeneration was observed in the CA3 or CA1 pyramidal cell layer (many positive cells with confluent fluorescence), the score related to this area was increased by 1. The scores obtained on NeuN and FJC sections were adjusted. The numbers resulting from these values were averaged per animal and used for statistical analysis.

### Immunohistochemical labelling

For immunofluorescence labelling, dorsal hippocampal sections from SHAM, SE, SE + VH-N412, SE + NT(8-13) and SE + DZP animals were pre-incubated for 1 H at RT in 0.12M PB containing 0.3% Triton X-100 and 3% BSA. Sections were then incubated overnight at 4°C in a solution containing the primary antibody: mouse anti-NeuN (1/1000; MAB377), or mouse anti-glial fibrillary acidic protein (GFAP; 1/1000, MAB360), both from Millipore, Darmstadt, Germany, or rabbit anti-ionized calcium binding adaptor molecule 1 (Iba1; 1/1000, Wako Pure Chemical Industries, Osaka, Japan), or rabbit anti-vesicular zinc transporter 3 (ZnT-3; 1/500; Synaptic System, Goettingen, Germany) diluted in PB 0.12 M containing 0.3% Triton X-100 and 3% BSA. After several rinses, sections were incubated for 2 H at RT in Alexa 488-conjugated anti-mouse or -rabbit IgG (1/200; Life Technologies) diluted in PB containing 3% BSA. Sections were then rinsed and coverslipped with Prolong Gold Anti-fading reagent (Life Technologies). In all immunohistochemical experiments, no labelling was detected when primary antibodies were omitted.

### Estimation of neuroinflammation

Astroglial (GFAP) and microglial (Iba1) reactivities were quantified by measuring labelling intensities on whole hippocampal sections from SHAM, SE, SE + VH-N412, SE + NT(8-13) and SE + DZP animals. The densitometric data were automatically calculated by ImageJ software. For each animal, the mean and corresponding SEM) intensity of labelling obtained from the total number of microscopic fields were calculated.

### Estimation of mossy fiber sprouting

Quantitative analysis was conducted to evaluate the number of mossy fiber terminals present in the DG IML from SHAM, SE, SE + VH-N412 and SE + NT(8-13) animals. These mossy fiber terminals revealed by ZnT-3 project aberrantly into this layer after KA-induced status epilepticus (Nadler et al., 1980; Sloviter et al., 2006; Epsztein et al., 2006). For this purpose, five Z-stacks of 3 confocal slices were acquired regularly along the IML for each animal, with a 40x objective and a numerical zoom 1,2. A region of interest (ROI) was outlined for each slice, and the terminals contained in the ROI were counted and averaged with the two other slices. The Cell Count plugin of *ImageJ* was used and counts were averaged to one final score per animal and mean numbers were used for statistical analysis.

### Behavioral analysis

#### Olfactory Tubing Maze (OTM)

The hippocampus-dependent learning and memory performance of control mice injected with saline solution (SHAM), SE and SE + VH-N412 mice were assessed 3 weeks after SE (12-week-old male mice). Behavioral assessment, based on the OTM, was carried out as we previously described (Roman et al., 2002; Girard et al., 2014). We have shown previously that the FVB/N mouse strain used in the present study performed very well in this task (Girard et al., 2016). Briefly, mice were trained to learn odor-reward associations. The mice placed on water restriction were submitted to the presentation of odors in testing chambers (TC). Two synthetic odors (strawberry and jasmine), coming from two distinct arms and randomly assigned to these arms, were simultaneously presented to the mice. Each of these 2 odors was arbitrarily associated either with a drop of water (the reward) or with a non-aversive but unpleasant sound, which were delivered depending on the arm chosen by the mice. The OTM is composed of 4 identical TC joined to each other forming a circular structure in which mice can move freely only in clockwise direction. The procedure is fully automated. The movement of the mice is detected by photoelectric cells. Odors, water, sound delivery, and the automatic doors separating the different TC were controlled by a microcomputer running with LabVIEW software (National Instruments, Colombes, France) that also automatically recorded the behavioral data. The learning procedure included 3 habituation days followed by 5 days of training, each daily sessions being composed of 12 trials/odor presentations corresponding to 3 clockwise laps/blocks of 4 trials. Two parameters were examined: the inter-trial interval (ITI) and the percentage of correct responses. The ITI was calculated as the time between the response to an odor presentation in one testing chamber and the response in the next chamber. The evolution of the ITI reflects the fact that, after a response to an odor, the animal must learn to backtrack to the testing chamber and to run to the next chamber waiting for the opening of the entrance door for the next trial. This constitutes a procedural aspect of the task. The percentage of correct responses was the ratio of the number of correct responses to the total number of odor presentations per session. This percentage was used to evaluate the effectiveness of the association between an odor and its reinforcement, a process that pertains to the hippocampus-dependent declarative memory subcategory.

#### Open field test

As described previously (Girard et al., 2016, 2014), the open field test was used to assess both exploratory behavior and spontaneous locomotor activity 4 weeks after SE and VH-N412 treatment. The mice were tested using an open field square made of white plastic with 50 x 50 cm surface area and 45 cm-high walls. Testing was carried out in a dimly illuminated room (40 lux). Monitoring was done by an automated tracking system using a video camera mounted above the apparatus (Viewpoint VideoTrack version 3.0, Champagne Au Mont D’or, France). The field was divided virtually into 2 regions of interest: a central area (20 x 20 cm) and a peripheral area. The animals were placed in the center of the field and 2 behavioral parameters were registered during 2 consecutive 5 min sessions: i) the percentage of time spent in the central part versus total time ii) the total traveled distance. The square was cleaned prior the test and between each animal with 70% ethanol.

### RT-qPCR and evaluation of cold shock protein mRNA levels

Total RNA was prepared from hippocampi of SHAM and VH-N412 treated mice for 4, 8 and 16 H and from rat mature hippocampal neuronal cultures treated with VH-N412 at 0.1, 1 and 10 μM using the Nucleospin RNA plus kit (Macherey Nagel, Allentown, PA USA). For the RT-qPCR experiments, all reagents, kits, equipment, and software were from Applied Biosystems. cDNA was synthesized from 500 ng of total RNA using the High-Capacity RNA-to-cDNA Kit. For real-time qPCR, 12.5 ng of cDNA were used. The samples were run in duplicate on 96-well plates and then analyzed with 7500 v2.0 software according to the manufacturer’s recommendations. All reactions were performed using TaqMan Fast Universal PCR Mix and TaqMan Assays probes (Table S4). The conditions of the thermal cycle were as follows: initial denaturation at 95°C for 40 cycles, denaturation at 95°C, and hybridization and extension at 60°C. Relative expression levels were determined according to the ΔΔCt (Ct: cycle threshold) method where the expression level of the mRNA of interest is given by 2^-ΔΔCT^, where ΔΔCT = ΔCt target mRNA - ΔCt reference mRNA (*Gapdh*, housekeeping gene) in the same sample.

### Immunoblot analysis of cold shock protein expression

Mouse brains were rapidly extracted, and the hippocampi from SHAM and VH-N412 treated animals were isolated and lysed in RIPA buffer (Sigma). After sonication, protein concentrations were determined using the Bio-Rad *DC*^TM^ protein assay kit (Bio-rad, Marnes-La-Coquette, France). Protein extracts (100 μg) were loaded and analyzed in 8% to 15% Tris-Glycine gels and subjected to western blotting (WB) with appropriate antibodies: rabbit anti-RBM3, (1/500, 14363-1-AP, Proteintech, Manchester, UK), CIRBP (1/1000, 10209-2-AP, Proteintech) and mouse anti-β-actin (1/5000, SC1615, Santa-Cruz, TX, USA). Briefly, proteins were transferred to a nitrocellulose membrane (GE Healthcare) and blocked 1 H with the appropriate blocking solution. Membranes were incubated overnight at 4°C with the primary antibodies and then with the corresponding horseradish peroxidase-conjugated secondary antibodies (Jackson Immunoresearch, West Grove, PA, USA). All membranes were revealed using ECL chemiluminescence kit according to the manufacturer’s instructions (GE Healthcare) and blots were analyzed with *ImageJ* software (NIH).

### Preparation of acute rat hippocampal slices, electrophysiological recordings and data analysis

Hippocampal slices (350 μm thick) were prepared from Sprague Dawley rats (n=3; 3-4 weeks old; Janvier Laboratories) and cut using a vibratome (Leica VT1200S) in an ice-cold oxygenated, modified ACSF, continuously aerated with 95% O_2_ and 5% CO_2_ and containing Glucose 11 mM, NaHCO_3_ 25 mM, NaCl 126 mM, KCl 3.5 mM, NaH_2_PO_4_ 1.2 mM, MgCl_2_ 1.3 mM and CaCl_2_ 2 mM. Slices were then incubated at RT for at least 1 hr in ACSF. Recordings were performed on hippocampal slices using multielectrode arrays (MEA). Slices were continuously perfused with the oxygenated ACSF at the rate of 3 mL/min with a peristaltic pump (MEA chamber volume: ∼1 mL). KA (300 nM) and VH-N412 (0.1, 1 and 10 μm) were added to the perfusion solution to assess the effects of VH-N412 compound on KA-induced increase of neuronal firing. Complete solution exchange in the MEA chamber was achieved 20 s after the switch of solutions. The perfusion liquid was continuously pre-heated at 37°C just before reaching the MEA chamber with a heated-perfusion cannula (PH01, MultiChannel Systems, Reutlingen, Germany). The temperature of the MEA chamber was maintained at 37 ± 0.1°C with a heating element located in the MEA amplifier headstage. The spike numbers per second recorded at each electrode were averaged for 30 s slots and normalized to the mean spikes rate value at t = 20-30 min (10 last minutes of KA exposure period). Individual data from independent experiments were then pooled and the mean values of the normalized spike rates (± SEM) were plotted as a function of time (before and after exposure to VH-N412). The control values (KA alone) were averaged from 3 rats, 3 slices and 18 electrodes. The dose response curves from the KA + VH-N412-treated slices were averaged from 3 rats, 4 slices and 25 electrodes.

### Primary cultures of rat hippocampal neurons, transfection and immunofluorescence

The primary hippocampal cells (mixed culture) were prepared from embryonic day 17-18 (E17 or E18) rats and cultured in Neurobasal supplemented with 2% B-27, 1% penicillin-streptomycin, and 0.3% glutamine in a humidified atmosphere containing 5% CO2 at 37°C. At 21 days *in vitro* (div) neurons displayed mature morphological and physiological features (Ivanov et al., 2009). Hippocampal cultures were immunostained for endogenous NTSR1 using goat polyclonal anti-NTSR1 (1/200, R20, Santa Cruz), dendritic marker using mouse anti-MAP2 (1/10000, Sigma), and for spine marker using rabbit anti-drebrin E/A (1/1000, Sigma). In some experiments, neurons were transiently co-transfected with RFP vector (red fluorescent protein) and with a plasmid construct encoding rat NTSR1 using lipofectamine 2000 reagent according to manufacturer’s protocol (ThermoFischer Scientific). Cells were fixed with 4% PFA in PB 0.12 M for 20 min at RT and immunostained for NTSR1 48 H following transfection. Cell coated coverslips were rinsed 3 times with PB 0.12 M and permeabilized in a blocking solution containing 3% BSA and 0.1% Triton X-100 diluted in PB 0.12 M for 30 min at RT. Cells were incubated overnight with primary antibodies diluted in 3% BSA blocking solution inside a humidity chamber at 4°C. Coverslips were washed 3 times for 5 min in PB 0.12 M, then incubated with corresponding secondary antibodies (Jackson ImmunoReasearch, Cambridgeshire, UK): donkey anti-goat IgG (H+L) highly cross-adsorbed AlexaFluor A594 (1/800), donkey anti-mouse IgG(H+L) highly cross-adsorbed AlexaFluor A 647 (1/800), and donkey anti-rabbit IgG (H+L) highly cross-adsorbed AlexaFluor A 488 (1/800) in PB 0.12 M containing 3% BSA at RT for 2 H and washed 3 times for 5 min each with PB 0.12 M. Nuclei were counterstained with 5 μg/mL DAPI at RT for 0.5 H. After 3 washes in PB 0.12 M, coverslips were rapidly rinsed 3 times in distilled water and let to dry before mounting on Superfrost glass slides using Fluoromount-G Mounting medium and stored at −20°C. Labelling specificity was assessed under the same conditions, by incubating some coverslips with non-transfected or transfected cells in a solution omitting the primary antibodies. In all cases, no overlap of antibodies was detected. Image acquisition was performed using Zen software and processed using Adobe Photoshop and *ImageJ* softwares.

### NMDA intoxication and drug treatment

For the glutamatergic agonist intoxication with NMDA or KA, cells were seeded at the density of 20,000 cells/well in 96 well-plates pre-coated with poly-L-lysine 10 μg/mL and cultured for 21 div. On day 21 of culture, the medium was removed, and fresh medium was added without or with test compounds (VH-N412, NTSR antagonists, BDNF (PAN-Biotech (Aidenbach, Germany) or Oestradiol (Sigma) 30 min before either NMDA (100 μM, 10 min) or KA (25 μM, 24 H) intoxication. BDNF (50 ng/mL) and Oestradiol (100 nM) were used as positive controls of neuroprotection against NMDA and KA induced injury respectively (Kajta and Lasoń, 2000; Mattson et al., 1995). After 24 H of NMDA or 48 H of KA intoxication, medium was removed and cells were washed twice in PB 0.12 M, followed by fixation with 4% PFA (Sigma) for 20 min at RT. Cells were permeabilized and non-specific sites were blocked with PB 0.12 M containing 0.1% saponin (Sigma) and 1% FCS (Invitrogen) and then incubated with mouse monoclonal anti-MAP-2. This antibody was revealed with Alexa Fluor 488 goat anti-mouse (Molecular probes, Eugene, OR, USA). Cell nuclei were labelled with Hoechst solution (Sigma). Neuronal death was assessed by counting the total number of neuronal cell bodies stained with Hoechst and MAP-2 and measuring the total dendrite length (all neurites stained with MAP-2) using Developer software (GE healthcare). For each condition, 10 pictures per well were taken using InCell 2000 (GE Healthcare) with 20x magnification. All the images were taken in the same conditions.

### Statistical analysis

Sample size and statistical power were determined according to Dell et al. (2002) and Festing and Altman (2002) using biostaTGV software (http://biostatgv.sentiweb.fr). All experiments were performed at least 3 times with different series of mice or independent cultures of rat hippocampal neurons. Student’s t-test was used to compare 2 groups. ANOVA analysis followed by Dunnett’s or Tukey’s *post hoc* test was used for multiple comparison. Behavioral data were analyzed with multivariate analyses of variance (MANOVAs) with repeated measures followed by Newman-Keuls post-hoc comparisons, using the SPSS/PC+ statistics 11.0 software (SPSS Inc., IL, USA) and selected ANOVAs were performed when necessary. All data were expressed as the mean ± SEM. Statistical significance was set at * *p*<0.05, ** *p*<0.01, and *** *p*<0.001.

## DISCUSSION

Research conducted in preclinical and clinical settings has demonstrated that mild to moderate hypothermia can provide neuroprotection in situations where there is an increased risk of neuronal death. These situations include sudden cardiac arrest followed by resuscitation, ischemic stroke, perinatal hypoxia/ischemia, traumatic brain injury (Kida et al., 2013; Andresen et al., 2015) and seizures (Sartorius and Berger, 1998; Schmitt et al., 2006; Gezalian, et al., 2015; Niquet et al., 2015a,b) in animal models and in humans (Karkar et al., 2002). Conversely, hyperthermia aggravates status epilepticus-induced epileptogenesis and neuronal loss in immature rats (Suchomelova et al., 2015). Hypothermia may open new therapeutic avenues for the treatment of epilepsy and for the prevention of its long-term consequences.

Intracerebral administration of NT or NT(8-13) analogues is associated with PIH (Bissette et al., 1976; Coquerel et al., 1988, 1986; Fanelli et al., 2015). When NT is administered peripherally, it is quickly metabolized by peptidases and has limited access through the BBB (McMahon et al., 2002; Gordon et al., 2003; Orwig et al., 2009; Boules et al., 2013). The conjugation of vector molecules can enhance the transport of active components across the BBB (Pardridge, 2001, 2003; Boer and Gaillard, 2007; Jones and Shusta, 2007; Pardridge, 2007; reviewed in Vlieghe and Khrestchatisky, 2013). Our objective was to generate “vectorized” forms of NT that cross the BBB and that display potent hypothermic properties, and to assess their potency in experimental epileptic conditions. For this purpose, we generated several conjugates that encompass peptides that target the LDLR and short active variants of NT. These were compared for their potential to bind LDLR on the one side and NTSR1 on the other using biophysical, cellular *in vitro* and *in vivo* approaches, and on their potential to induce hypothermia in different species, including mice, rats and pigs. Based on this comparison, we selected the VH-N412 conjugate for further studies and evaluated its neuroprotective properties in a mouse model of temporal lobe epilepsy. We showed that this compound reduced epileptic seizures, neurodegeneration, neuroinflammation, mossy fiber sprouting and preserved learning and memory skills in the epileptic mice. We showed that the target receptor NTSR1, was expressed in hippocampal pyramidal neurons *in vitro* and *in vivo*, in cell bodies, dendrites and spines. Besides the neuroprotective hypothermia effects induced *in vivo* with VH-N412, we showed in cultured hippocampal neurons that this compound also displayed temperature-independent neuroprotective properties.

### LDLR binding peptides conjugated to NT induce potent hypothermia in naïve mice when administered systemically

We coupled LDLR-binding peptides to NT and optimization and SAR evaluations were performed to facilitate synthesis together with improving biological properties of the conjugate, namely LDLR- and NTR1-binding, metabolic stability, BBB-crossing and, in the end, central hypothermic potential after systemic administration in mice. Starting with the VH-N21 and VH-N41 conjugates encompassing respectively the VH0445 and VH04129 peptides conjugated to the full-length NT tridecapeptide, we selected the VH-N412 conjugate encompassing the VH04129 and NT(8-13) peptides, to generate a low molecular weight conjugate (1929 kDa) that displayed the most potent and sustained hypothermic potential in naïve mice at low dose (ED50 *ca.* 0.8 mg/kg, corresponding to 0.7 mg/kg eq. NT). Consistent with previous reports with other LDLR-binding VH445 peptide analogues (Jacquot et al., 2016; David et al., 2018; Varini et al., 2019) the VH04129 peptide retained its binding potential to LDLR when associated to the NT(8-13) peptide (63.8 nM), compared to the free VH4129 (64.8 nM). On the other hand, binding of the NT(8-13) fragment to the NTSR-1, which was previously shown to mediate central hypothermia (Pettibone et al., 2002; Remaury et al., 2002; Mechanic et al., 2009), was also retained when associated to VH4129. Interestingly, the VH4129 peptide analogue was evaluated based on its optimal LDLR-binding profile, with moderate affinity (similar to the VH445 analogue) but with faster on-rate and off-rate than VH445 (Jacquot et al., 2016), endowing this analogue with a better binding/release profile for BBB-crossing. Accordingly, the BBB influx rate (K_in_) of the VH-N412 compound was 3-fold higher than the initial VH-N21 compound encompassing the VH445 peptide, and 10-fold higher than the native NT peptide. Although only sparse data exists evaluating this parameter for NT analogues (Banks et al., 1995), this result confirms the optimized brain uptake potential of the VH-N412 compound. Finally, because the primary cleavage site of the NT(8-13) peptide in plasma is at the N-terminal Arg-Arg site (Orwig et al., 2009), conjugation of the VH4129 peptide probably protects to some extent from aminopeptidases. This was confirmed by the extended metabolic resistance of VH-N412 *in vitro* in both mouse and human blood, with more than 1 H half-life vs. 44 min with the VH445, previously found to be less stable than the optimized VH4129 analogue (Jacquot et al., 2016), and only 9 min for the native NT. This was also consistent with the rapid enzymatic proteolysis of endogenous peptides and generally small linear peptides containing only natural amino-acids, highlighting the advantage of using a fully optimized LDLR-binding peptide (Foltz et al., 2010). The *in situ* brain perfusion results showed clearly that the VH-N21, and more so the VHN-412, displayed higher brain uptake in mice than non-conjugated NT. However, we cannot exclude that the hypothermia we observed with the different conjugates is related in part to increased stability conferred to NT.

### VH-N412 induces potent PIH, and attenuates seizures, neurodegeneration and neuroinflammation in the KA model of epilepsy

Following induction of SE with KA, we show that the VH-N412 compound elicited rapid hypothermia that was associated with anticonvulsant effects. In particular, we observed potent neuroprotection, reduced inflammation in the hippocampus, reduced sprouting of the DG mossy fibers and preserved learning and memory skills in the epileptic mice treated with VH-N412. To our knowledge, these results are the first to show a significant impact of PIH in an epileptic mouse model. Hypothermia appears to alleviate seizure intensity very rapidly, as early as 30 min after VH-N412 administration, and as efficiently as DZP, one of the most efficient anti-seizure drugs that interestingly, also induces hypothermia (Irvine, 1966). Several studies show that NT or stabilized NT analogs display anticonvulsant properties while hypothermia was apparently not evaluated in these studies (Green et al., 2010; Lee et al., 2009; Robertson et al., 2011; Clynen et al., 2014). We thus cannot exclude that to some extent, our NT-peptide conjugates also reduce seizures independently of hypothermia.

The neuroprotective effects we observed are likely due to reduced seizure burden but also possibly to reduced brain metabolism during the phases that follow seizures. Such neuroprotective effects are comparable to those discussed below in several studies of acute brain damage in rodent models of hypoxia-ischemia, traumatic brain injury (TBI) and intracerebral hemorrhage (ICH) using different NT analogs. Among these, NT77 was modified from NT at amino-acid positions located in the hexapeptide of neurotensin NT[8-13]) (Boules et al., 2001). NT77 crossed the BBB, induced hypothermia (Gordon et al., 2003), reduced oxidative stress in the rat hippocampus (Katz et al., 2004a) and improved neurologic outcome after asphyxial cardiac arrest (Katz et al., 2004b). ABS-201, also known as HPI-201, another synthetic analogue of NT[8-13] has been described as a second-generation high affinity NTSR1 agonist that exhibited BBB permeability, and effectively induced PIH in a number of experimental paradigms. In a focal ischemic model of adult mice ABS-201 induced recovery of sensory motor function (Choi et al., 2012). It also promoted the integrity of the BBB and the neurovascular unit (Zhao et al., 2020). HPI-201 greatly enhanced the efficiency and efficacy of conventional physical cooling in a rat model of ischemic stroke (Lee et al., 2016a). In the same model in mice and oxygen glucose deprivation in cortical neuronal cultures, HPI-201 displayed anti-inflammatory effects (Lee et al., 2016b). HPI-201-induced hypothermia saved neurons and endothelial cells inside the ischemic core in mice (Jiang et al., 2017). In a ventricular fibrillation cardiac arrest (VFCA) rat model, ABS 201 induced therapeutic hypothermia, ameliorated post-resuscitation myocardial-neurological dysfunction, and prolonged survival duration (Li et al., 2019). HPI-201 was effective in reducing neuronal and BBB damage, attenuating inflammatory response and detrimental cellular signaling, and promoting functional recovery after TBI in the developing rat brain (Gu et al., 2015). HPI-201 also protected the brain from ICH injury in mice (Wei et al., 2013). Another second-generation NTSR1 agonist, HPI-363 elicited hypothermia and protective effects, improving sensorimotor functional recovery in a TBI model in adult rat brain (Lee et al., 2014) and induced beneficial effects on chronically developed post-stroke neuropsychological disorders, in the pre-frontal cortex of mice (Zhong et al., 2020).

### NT receptors potentially involved in VH-N412-induced PIH in the KA model of epilepsy

NT-producing neurons and their projections are widely distributed in the brain, including the anterior hypothalamus (Schroeder and Leinninger, 2018) involved in thermoregulation. It is likely that NT-induced hypothermia acts via similar processes in the epilepsy model used in this study and in the different models of acute brain damage mentioned above, reducing both excitotoxicity and neuroinflammation in different vulnerable regions of the CNS. While the mechanisms leading to hypothermia remain largely unknown, the NT analogs are known to modulate the activity of NT receptors. mRNA encoding the high affinity NTSR1 was detected essentially in neurons in many hypothalamic regions, including the preoptic, anterior, periventricular, ventromedial and arcuate nuclei (Alexander and Leeman, 1998; Nicot et al., 1994; Woodworth et al., 2018). In contrast, transcripts encoding the low affinity NTSR2 were present essentially in astrocytes (Woodworth et al., 2018), as we also recently demonstrated in the hippocampus (Kyriatzis et al., 2021). Studies with NTSR1^−/−^ and NTSR2^−/−^ mice and the absence of hypothermia effects of NTSR2-selective analogs suggested that in the different pathophysiological conditions evoked above, hypothermia is mediated by NTSR1 (Pettibone et al., 2002; Remaury et al., 2002; Mechanic et al., 2009; Boules et al., 2010). However, this remains controversial since receptor knockdown using antisense oligodeoxynucleotides in adult mice point to an involvement of NTSR2 in hypothermia (Dubuc et al., 1999). More recently, it appeared that activation of both NTSR1 and NTSR2 was required for a full hypothermic response (Tabarean, 2020).

### VH-N412 preserves learning and memory abilities, exploratory and normal locomotor activity in the KA model of epilepsy

Using two behavioral tasks, we report here that treatment of mice with VH-N412 after SE preserves learning and memory abilities as well as exploratory and general spontaneous motor activity. Hippocampus-dependent learning and memory performance was assessed using the OTM, in which mice are expected to make odor-reward associations in darkness, without visual cues (Roman et al., 2002). This test was selected in regard to the sensorial abilities of the FVB/N strain that are homozygous for the Pde6brd1 allele with an early onset retinal degeneration and blindness (Brown and Wong, 2007). Consequently, they are deeply impaired in spatial tasks such as the Morris water maze (Brown and Wong, 2007; Owen et al., 1997; Pugh et al., 2004; Royle et al., 1999; Võikar et al., 2001) or radial maze (Mineur and Crusio, 2002). In the OTM, it has already been demonstrated that albino mouse strains with reduced visual abilities such as BALB/C or CD1 mice are able to learn the task as well as or better than mouse strains with functional vision (Restivo et al., 2006; Roman et al., 2002). FVB/N mice reached a level of 80 ± 5 % of correct responses from the fourth 12-trial training session. It has been shown that bilateral excitotoxic lesions of the hippocampus induced by injections of ibotenic acid, another glutamate agonist, led to learning and memory impairments of BALB/C mice in the OTM (Nivet et al., 2011). Using the FVB/N mice similar learning and memory deficits were observed after injections of KA and consecutive seizures. In SE mice treated with VH-N412, we observed similar learning and memory performance compared to control mice. Similarly, behavioral activity was preserved in spontaneous locomotor activity using the open field test. SE and related hippocampal excitotoxic lesions are known to induce hyperactivity in rats (dos Santos et al., 2000; Kubová et al., 2004) and mice (Chen et al., 2002; Müller et al., 2009), which can be easily highlighted using the open field test. As expected, our experiments demonstrated a strong hyperactivity in FVB/N mice four weeks after KA injections and SE, that was not observed in animals treated with VH-N412.

### Cellular and molecular mechanisms potentially modulated by VH-N412 in the KA model of epilepsy

NT, its analogs, agonists and antagonists have been shown to modulate both GABAergic and glutamatergic activity (Ferraro et al., 2008; Li et al., 2008). We investigated whether VH-N412 could modulate hippocampal neuronal hyperactivity induced by KA using MAE on acute hippocampal slices, but this was not the case. These observations suggest that VH-N412 does not modulate neuronal hyperactivity *in vivo*, at least in the hippocampus. Modulators of the NT system may regulate neuroprotective pathways that are probably common to the different models of brain pathology evoked above. These models display neuronal excitotoxicity (reviewed in (Mattson, 2017), neuroinflammation, as shown by pro-inflammatory glial markers such as astrocytic GFAP or microglial Iba1, endothelial and BBB damage (Kyriatzis et al., 2021, 2024). It has been shown in different models that PIH increased BDNF and vascular endothelial growth factor (VEGF), reduced the pro-apoptotic caspase-3 activation, BAX, and MMP-9 expression, decreased expression of inflammatory factors including monocyte chemoattractant protein-1 (MCP-1) and macrophage inflammatory protein-1α (MIP-1α), two key chemokines in the regulation of microglia activation and infiltration. Moreover, up-regulation of the M1 microglia markers IL-12, IL-23, inducible nitric oxide synthase (iNOS), of tumor necrosis factor-α (TNF-α), interleukin-1β (IL-1β) and IL-6 was also shown to be decreased or prevented. Meanwhile, TH treatments increased the expression of M2 type reactive anti-inflammatory factors including IL-10, Fizz1, Ym1, and arginase-1 and the expression of the anti-apoptotic Bcl-2 (Choi et al., 2012; Wei et al., 2013; Lee et al., 2014; Gu et al., 2015; Lee et al., 2016; Jiang et al., 2017; Zhao et al., 2020).

### Cold shock proteins RBM3 and CIRBP are involved in VH-N412-induced PIH in the KA model of epilepsy

Among the proteins known to be modulated by hypothermia are cold-shock proteins such as RBM3 and CIRBP. They are known to escape translational repression, and are involved in diverse physiological and pathological processes, including hibernation, circadian rhythm, inflammation, neural plasticity, and they modulate neurodegeneration in relation with body temperature (Williams et al., 2005; Smart et al., 2007; Chip et al., 2011; Tong et al., 2013; reviewed in Zhu et al., 2016; Ávila-Gómez et al., 2020).

Following VH-N412 administration that induces hypothermia in naïve mice, we observed significant increase of mRNA encoding RBM3 and CIRBP in hippocampus relative to control at 16 and 8 H time points respectively, correlated with RBM3 and CIRBP protein increase. In agreement with previous studies (Peretti et al., 2015), the absence of modulation of RBM3 and CIRBP mRNA we observed in neurons cultured at 37°C and treated with different concentrations of VH-N412, suggests that it is indeed hypothermia that induced expression of these cold shock proteins in our epileptic model. Induction of these cold shock proteins in epileptic mice treated with VH-N412 is probably associated with the observed neuroprotection as shown in other experimental paradigms (Peretti et al., 2015).

### VH-N412 has intrinsic neuroprotective effects in cultured hippocampal neurons, independent of its potential to induce hypothermia

The hippocampus is known as a highly vulnerable structure of the CNS, with rapid loss of principle cells including pyramidal cells and interneurons (reviewed in Houser, 2014). Remarkably, these neurons were protected by VH-N412 administration in our epilepsy model. Two processes may be involved in such neuroprotection: i) either hypothermia with a global reduction of cell metabolism, oxidative stress and excitotoxic processes, with concomitant reduction of deleterious neuroinflammation; ii) intrinsic neuroprotective effects of VH-N412 that could act in synergy with hypothermia. These 2 processes are not exclusive and it is possible that the conjugate causes hypothermia and has favorable effects on the sequelae of SE. Further experiments are warranted where one could prevent pharmacological hypothermia by warming up the animals undergoing SE and administered with VH-N412, to evaluate the intrinsic neuroprotective effects of the conjugate *in vivo*.

Our immunohistochemistry experiments on mouse brain sections showed that hippocampal neurons expressed the NTSR1 receptor, that is located not only in the neuronal cell bodies, but also in proximal dendrites and in spines. This was confirmed in cultured hippocampal neurons expressing either the endogenous or exogenous NTSR1. These observations lead us to question whether VH-N412 displayed neuroprotective effects on pyramidal neurons in culture. These were challenged with two excitotoxic agents, NMDA and KA, that induce excitotoxic neuronal death via their respective receptors. We showed that VH-N412 did induce neuroprotection as efficiently as molecules well-known for neuroprotection, BDNF and Oestradiol respectively (Mattson et al., 1995; Kajta and Lasoń, 2000). This is in agreement with data showing that the neurotoxic effects of methamphetamine on dopaminergic terminals and apoptosis of striatal neurons were attenuated by the NTSR1 agonist PD149163, independently of hypothermia (Liu et al., 2017). In another study, exogenous treatment with the PD149163 NTSR1 agonist exerted neuroprotection in HFD-induced pre-diabetic rats and Alzheimer models. It was hypothesized that the treatment with PD149163 attenuates hippocampal pathologies and synaptic dysfunction, eventually restoring cognition through hippocampal NTSR1 signaling. In particular, it was shown that the NTSR1 agonist PD149163 increased dendritic spines and reduced hippocampal microglial cells (Saiyasit et al., 2021). Finally, as mentioned above, NTSR agonists induced neuroprotection in a number of experimental conditions including global ischemia, stroke, TBI etc. At this point it is difficult to determine the direct contribution of hypothermia compared to intrinsic neuroprotective effects of these agonists. In contrast, other studies point to the deleterious effects of NTSR agonists on glutamatergic excitotoxicity both *in vitro* and *in vivo*. NT enhanced glutamate-induced excitotoxicity through NTSR1 activation in both mesencephalic and cortical neurons (Antonelli et al., 2004, 2002). NT increased glutamate release in some brain regions (Sanz et al., 1993; Ferraro et al., 1995) and it has been shown that in addition to enhancing glutamate release, NT can modify the function of glutamate receptors *in vitro* and *in vivo* (Antonelli et al., 2004; Ferraro et al., 2011, 2008).

## CONCLUSION

In all, our results suggest that if indeed VH-N412 exerts neuroprotective effects via hypothermia in our epileptic model, there is some *in vitro* evidence that this molecule exhibits neuroprotective effects independently from hypothermia. This hypothesis will need to be challenged further *in vitro* and in other models of brain diseases *in vivo*. In the present article, we essentially focused on epilepsy and acute neurotoxicity models and showed the beneficial effects of a vectorized active peptide such as NT following systemic administration. The CNS is endowed with many devastating pathologies including neurodegenerative diseases. The latter could benefit from new drugs, some of them in development, including biologics such as oligonucleotides, RNAi, proteins or therapeutic antibodies, at the condition they reach the diseased CNS. Thus, our results emphasize the potential of drug delivery approaches based on targeting peptide or antibody fragment moieties to bring conjugated biologics across the BBB into the diseased CNS, following systemic administration of conjugates.

## ACKNOWLEDGEMENTS

Financial support was provided by the French National Agency for Research (VECtoBrain ANR-09-BIOT-015-01 to VECT-HORUS, VEC2Brain ANR-13-RPIB-0010-01 to MK and NANOVECTOR ANR-15-CE18-0010-03 to MK), by Aix Marseille Université and by the CNRS. We thank the animal facility of the Faculty of Pharmacy of Paris-Cité Université (US25 Inserm, UAR3612 CNRS) for hosting the animals for the *in situ* brain perfusion experiments.

