## Supplementary material for "A peptide-neurotensin conjugate that crosses the blood-brain barrier induces pharmacological hypothermia associated with anticonvulsant, neuroprotective and anti-inflammatory properties following status epilepticus in mice": Table S1

Table S1. Analytical characterization of Peptide-NT conjugates

| Compound |  | Peptide-vector |  | Neurotensin |  | Coupling strategy |  |  | Yield (%) |  | MW (g/mol)  observed/calculated |
| --- | --- | --- | --- | --- | --- | --- | --- | --- | --- | --- | --- |
| **VH-N21** |  | **VH445** |  | Full length NT(1-13) |  | Thiol-Mal on NT(Lys6) |  |  | 39% |  | 2958.52/2958.45 |
| **VH-N41** |  | **VH4129** |  | Full length NT(1-13) |  | Thiol-Mal on NT(Lys6) |  |  | 40% |  | 2998.50/2998.57 |
| **VH-N42** |  | **VH4129** |  | NT(2-13) |  | Thiol-Mal on NT(Lys6) |  |  | 43% |  | 2929.47/2929.69 |
| **VH-N43** |  | **VH4129** |  | NT(6-13) |  | Thiol-Mal on NT(Lys6) |  |  | 46% |  | 2410.24/2410.38 |
| **VH-N44** |  | **VH4129** |  | Lys-NT(8-13) |  | Thiol-Mal on Lys |  |  | 42% |  | 2313.19/2313.24 |
| **VH-N46** |  | **VH4129** |  | GGG-NT(6-13) |  | Tandem (linear) |  |  | 4% |  | 2212.17/2212.10 |
| **VH-N49** |  | **VH4129** |  | Ahx-NT(8-13) |  | Tandem (linear) |  |  | 21% |  | 1929.04/1929.00 |
| **VH-N412** |  | **VH4129** |  | PEG6-NT(8-13) |  | Tandem (linear) |  |  | 17% |  | 2151.15/2151.50 |

Table S1 : Ferhat et al.
