## Supplementary material for "A peptide-neurotensin conjugate that crosses the blood-brain barrier induces pharmacological hypothermia associated with anticonvulsant, neuroprotective and anti-inflammatory properties following status epilepticus in mice": Table S2

Table S2. Hypothermic response to IV administration of Peptide-NT conjugates in mice

| Compound |  | Dose  (mg/kg eq. NT) |  | DBT (°C) ± sem |  | T_max_ (min) |  | BT (°C) ± sem  at T_max_ ^(1)^ |
| --- | --- | --- | --- | --- | --- | --- | --- | --- |
| **VH-N21** |  | 0.5 |  | -0.9 ± 0.3 |  | 15 |  | 37.3 ± 0.3 |
|  |  | 10 |  | -3.7 ± 0.3 |  | 60 |  | 34.2 ± 0.5 |
| **VH-N41** |  | 0.5 |  | -2.3 ± 0.3 |  | 30 |  | 36.2 ± 0.3 |
|  |  | 10 |  | -6.8 ± 0.2 |  | 60 |  | 31.6 ± 0.2 |
| **VH-N42** |  | 0.5 |  | -2.5 ± 0.3 |  | 30 |  | 36.1 ± 0.3 |
|  |  | 10 |  | ND |  | ND |  | ND |
| **VH-N43** |  | 0.5 |  | -2.9 ± 0.4 |  | 30 |  | 35.4 ± 0.4 |
|  |  | 10 |  | ND |  | ND |  | ND |
| **VH-N44** |  | 0.5 |  | -2.3 ± 0.6 |  | 30 |  | 36.7 ± 0.5 |
|  |  | 10 |  | ND |  | ND |  | ND |
| **VH-N46** |  | 0.5 |  | -1.2 ± 0.5 |  | 15 |  | 37.3 ± 0.4 |
|  |  | 10 |  | -5.6 ± 0.8 |  | 60 |  | 33.0 ± 0.7 |
| **VH-N49** |  | 0.5 |  | -2.6 ± 0.3 |  | 30 |  | 35.7 ± 0.4 |
|  |  | 10 |  | -6.8 ± 0.3 |  | 90 |  | 31.4 ± 0.3 |
| **VH-N412** |  | 0.5 |  | -2.7 ± 0.4 |  | 30 |  | 35.3 ± 0.3 |
|  |  | 10 |  | -6.7 ± 0.6 |  | 60 |  | 32.5 ± 0.6 |
| ND not determined; DBT: maximal change in Body Temperature; T_max_: time post-injection to maximal change in body temperature; (1) Baseline temperature in mice was 38-39°C.  Table S2 : Ferhat et al. | | | | | | | | |
