## Supplementary material for "A peptide-neurotensin conjugate that crosses the blood-brain barrier induces pharmacological hypothermia associated with anticonvulsant, neuroprotective and anti-inflammatory properties following status epilepticus in mice": Table S3

Table S3. Comparison of mean body temperature changes between SHAM, SE, SE + VH-N412, SE + NT (8-13) and SE + DZP

Mean body temperature changes (°C) ± SEM	*P*	Tukey’s test

```
-30 min

SHAM vs SE                38,64±0,20 vs 38,73±0,20      ns
SHAM vs SE+VH-N412        38,64±0,20 vs 38,58±0,22      ns
SHAM vs SE+NT(8-13)       38,64±0,20 vs 39,11±0,29      ns
SHAM vs SE+DZP            38,64±0,20 vs 39,24±0,28      ns
SE vs SE+VH-N412          38,73±0,21 vs 38,58±0,22      ns
SE vs SE+NT(8-13)         38,73±0,21 vs 39,11±0,29      ns
SE vs SE+DZP              38,73±0,21 vs 39,24±0,28      ns
SE+VH-N412 vs SE+NT(8-13) 38,58±0,22 vs 39,11±0,29      ns
SE+VH-N412 vs SE+DZP      38,73±0,21 vs 39,24±0,28      ns

0 min

SHAM vs SE                38,64±0,25 vs 39,08±0,40      ns
SHAM vs SE+VH-N412        38,64±0,25 vs 39,23±0,45      ns
SHAM vs SE+NT(8-13)       38,64±0,25 vs 40,00±0,28      ns
SHAM vs SE+DZP            38,64±0,25 vs 40,30±0,31      ns
SE vs SE+VH-N412          39,08±0,40 vs 39,23±0,45      ns
SE vs SE+NT(8-13)         39,08±0,40 vs 40,00±0,28      ns
SE vs SE+DZP              39,08±0,40 vs 40,30±0,31      ns
SE+VH-N412 vs SE+NT(8-13) 39,23±0,45 vs 40,00±0,28      ns
SE+VH-N412 vs SE+DZP      39,23±0,45 vs 40,30±0,31      ns

30 min

SHAM vs SE                38,38±0,15 vs 38,62+0,37      ns
SHAM vs SE+VH-N412        38,38±0,15 vs 36,50±0,34      ns
SHAM vs SE+NT(8-13)       38,38±0,15 vs 38,58±0,39      ns
SHAM vs SE+DZP            38,38±0,15 vs 36,10±0,44      ns
SE vs SE+VH-N412          38,62±0,37 vs 36,50±0,34   <0,01
SE vs SE+NT(8-13)         38,62±0,37 vs 38,58±0,39      ns
SE vs SE+DZP              38,62±0,37 vs 36,10±0,44   <0,01
SE+VH-N412 vs SE+NT(8-13) 38,62±0,37 vs 38,58±0,39   <0,01
SE+VH-N412 vs SE+DZP      38,62+0,37 vs 36,10+0,44      ns

60 min

SHAM vs SE                38,12±0,30 vs 39,43±0,30      ns
SHAM vs SE+VH-N412        38,12±0,30 vs 35,27±0,32   <0,01
SHAM vs SE+NT(8-13)       38,12+0,30 vs 38,30+0,54      ns
SHAM vs SE+DZP            38,12+0,30 vs 34,80+0,41   <0,01


SE vs SE+VH-N412          39,43±0,30 vs 35,27±0,32   <0,01
SE vs SE+NT(8-13)         39,43±0,30 vs 38,30±0,54      ns
SE vs SE+DZP              39,43±0,30 vs 34,80±0,41   <0,01
SE+VH-N412 vs SE+NT(8-13) 35,27±0,32 vs 38,30±0,54   <0,01
SE+VH-N412 vs SE+DZP      35,27±0,32 vs 34,80±0,41      ns

90 min

SHAM vs SE                37,88±0,41 vs 39,08±0,23      ns
SHAM vs SE+VH-N412        37,88±0,41 vs 35,07±0,41   <0,01
SHAM vs SE+NT(8-13)       37,88±0,41 vs 37,78±0,49      ns
SHAM vs SE+DZP            37,88±0,41 vs 34,38±0,60   <0,01
SE vs SE+VH-N412          39,00±0,23 vs 35,07±0,41   <0,01
SE vs SE+NT(8-13)         39,08±0,23 vs 37,78±0,49      ns
SE vs SE+DZP              39,08±0,23 vs 34,38±0,60   <0,01
SE+VH-N412 vs SE+NT(8-13) 35,07±0,41 vs 37,78±0,49   <0,01
SE+VH-N412 vs SE+DZP      35,07+0,41 vs 34,38±0,60      ns

120 min

SHAM vs SE                37,78±0,44 vs 37,78±0,44      ns
SHAM vs SE+VH-N412        37,78±0,44 vs 34,74±0,43   <0,01
SHAM vs SE+NT(8-13)       37,78±0,44 vs 37,53±0,39      ns
SHAM vs SE+DZP            37,78±0,44 vs 34,32±0,86   <0,01
SE vs SE+VH-N412          37,78±0,44 vs 34,74±0,43   <0,01
SE vs SE+NT (8-13)        37,78±0,44 vs 37,53±0,39      ns
SE vs SE+DZP              37,78±0,44 vs 34,32±0,86   <0,01
SE+VH-N412 vs SE+NT(8-13) 34,74±0,43 vs 37,53±0,39   <0,01
SE+VH-N412 vs SE+DZP      34,74+0,43 vs 34,32±0,86      ns

150 min

SHAM vs SE                37,92±0,30 vs 38,44±0,22      ns
SHAM vs SE+VH-N412        37,92±0,30 vs 35,10±0,42   <0,01
SHAM vs SE+NT(8-13)       37,92±0,30 vs 37,40±0,37      ns
SHAM vs SE+DZP            37,92±0,30 vs 34,72±0,89   <0,01
SE vs SE+VH-N412          38,44±0,22 vs 35,10±0,42   <0,01
SE vs SE+NT (8-13)        38,44±0,22 vs 37,40±0,37      ns
SE vs SE+DZP              38,44±0,22 vs 34,72±0,89   <0,01
SE+VH-N412 vs SE+NT(8-13) 35,10±0,42 vs 37,40±0,37   <0,01
SE+VH-N412 vs SE+DZP      35,10±0,42 vs 34,72±0,89      ns
```

SHAM (n=5 mice) ; SE (n=16 mice) ; VH-N412 (n=20) ; SE+NT(8-13)(n=14) ; SE+DZP (n=5 mice) ; ns, not significant.

Table S3 : Ferhat et al.
