## Supplementary material for "A peptide-neurotensin conjugate that crosses the blood-brain barrier induces pharmacological hypothermia associated with anticonvulsant, neuroprotective and anti-inflammatory properties following status epilepticus in mice": Table S4

Table S4. Comparison of mean seizure intensity score between SHAM, SE, SE + VH-N412, SE + NT (8-13) and SE + DZP

Mean seizure intensity score ± SEM	*P*	Tukey’s test

```
-30 min

SHAM vs SE                0,00±0,00 vs 5,19±0,10   <0,01
SHAM vs SE+VH-N412        0,00±0,00 vs 5,30±0,11   <0,01
SHAM vs SE+NT(8-13)       0,00±0,00 vs 5,00±0,00   <0,01
SHAM vs SE+DZP            0,00±0,00 vs 5,00±0,00   <0,01
SE vs SE+VH-N412          5,19±0,10 vs 5,30±0,11      ns
SE vs SE+NT(8-13)         5,19±0,10 vs 5,00±0,00      ns
SE vs SE+DZP              5,19±0,10 vs 5,00±0,00      ns
SE+VH-N412 vs SE+NT(8-13) 5,30±0,11 vs 5,00±0,00      ns
SE+VH-N412 vs SE+DZP      5,30±0,11 vs 5,00±0,00      ns

0 min

SHAM vs SE                0,00±0,00 vs 5,25±0,17   <0,01
SHAM vs SE+VH-N412        0,00±0,00 vs 5,40±0,11   <0,01
SHAM vs SE+NT(8-13)       0,00±0,00 vs 5,14±0,10   <0,01
SHAM vs SE+DZP            0,00±0,00 vs 5,00±0,00   <0,01
SE vs SE+VH-N412          5,25±0,17 vs 5,40±0,11      ns
SE vs SE+NT(8-13)         5,25±0,17 vs 5,14±0,10      ns
SE vs SE+DZP              5,25±0,17 vs 5,00±0,00      ns
SE+VH-N412 vs SE+NT(8-13) 5,40±0,11 vs 5,14±0,10      ns
SE+VH-N412 vs SE+DZP      5,40±0,11 vs 5,00±0,00      ns


30 min

SHAM vs SE                0,00±0,00 vs 5,38±0,15   <0,01
SHAM vs SE+VH-N412        0,00±0,00 vs 1,97±0,36      ns
SHAM vs SE+NT(8-13)       0,00±0,00 vs 4,71±0,46   <0,01
SHAM vs SE+DZP            0,00±0,00 vs 0,80±0,80      ns
SE vs SE+VH-N412          5,38±0,15 vs 1,97±0,36   <0,01
SE vs SE+NT(8-13)         5,38±0,15 vs 4,71±0,46      ns
SE vs SE+DZP              5,38±0,15 vs 0,80±0,80   <0,01
SE+VH-N412 vs SE+NT(8-13) 1,97±0,36 vs 4,71±0,46   <0,01
SE+VH-N412 vs SE+DZP      1,97±0,36 vs 0,80±0,80      ns

60 min

SHAM vs SE                0,00±0,00 vs 5,37±0,15   <0,01
SHAM vs SE+VH-N412        0,00±0,00 vs 1,57±0,32      ns
SHAM vs SE+NT(8-13)       0,00±0,00 vs 4,85±0,25   <0,01
SHAM vs SE+DZP            0,00±0,00 vs 0,80±0,80      ns
SE vs SE+VH-N412          5,37±0,15 vs 1,57±0,32   <0,01


SE vs SE+NT(8-13)         5,37±0,15 vs 4,85±0,25      ns
SE vs SE+DZP              5,37±0,15 vs 0,80±0,80   <0,01
SE+VH-N412 vs SE+NT(8-13) 1,57±0,32 vs 4,85±0,25   <0,01
SE+VH-N412 vs SE+DZP      1,57±0,32 vs 0,80±0,80      ns

90 min

SHAM vs SE                0,00±0,00 vs 5,43±0,13   <0,01
SHAM vs SE+VH-N412        0,00±0,00 vs 1,36±0,34      ns
SHAM vs SE+NT(8-13)       0,00±0,00 vs 4,92±0,25   <0,01
SHAM vs SE+DZP            0,00±0,00 vs 0,40±0,40      ns
SE vs SE+VH-N412          5,43±0,13 vs 1,36±0,34   <0,01
SE vs SE+NT(8-13)         5,43±0,13 vs 4,92±0,25      ns
SE vs SE+DZP              5,43±0,13 vs 0,40±0,40   <0,01
SE+VH-N412 vs SE+NT(8-13) 1,36±0,34 vs 4,92±0,25   <0,01
SE+VH-N412 vs SE+DZP      1,36±0,34 vs 0,40±0,40      ns

120 min

SHAM vs SE                0,00±0,00 vs 5,18±0,14   <0,01
SHAM vs SE+VH-N412        0,00±0,00 vs 1,10±0,27      ns
SHAM vs SE+NT(8-13)       0,00±0,00 vs 4,85±0,23   <0,01
SHAM vs SE+DZP            0,00±0,00 vs 0,00±0,00      ns
SE vs SE+VH-N412          5,18±0,14 vs 1,10±0,27      <0,01
SE vs SE+NT(8-13)         5,18±0,14 vs 4,85±0,23      ns
SE vs SE+DZP              5,18±0,14 vs 0,00±0,00   <0,01
SE+VH-N412 vs SE+NT(8-13) 1,10±0,27 vs 4,85±0,23   <0,01
SE+VH-N412 vs SE+DZP      1,10±0,27 vs 0,00±0,00      ns

150 min

SHAM vs SE                0,00±0,00 vs 5,25±0,11   <0,01
SHAM vs SE+VH-N412        0,00±0,00 vs 1,26±0,27      ns
SHAM vs SE+NT(8-13)       0,00±0,00 vs 4,85±0,23   <0,01
SHAM vs SE+DZP            0,00±0,00 vs 0,00±0,00      ns
SE vs SE+VH-N412          5,25±0,11 vs 1,26±0,27   <0,01
SE vs SE+NT(8-13)         5,25±0,11 vs 4,85±0,23      ns
SE vs SE+DZP              5,25±0,11 vs 0,00±0,00   <0,01
SE+VH-N412 vs SE+NT(8-13) 1,26±0,27 vs 4,85±0,23   <0,01
SE+VH-N412 vs SE+DZP      1,26±0,27 vs 0,00±0,00      ns
```

SHAM (n=5 mice) ; SE (n=16 mice) ; VH-N412 (n=20) ; SE+NT(8-13)(n=14) ; SE+DZP (n=5 mice) ; ns, not significant.

Table S4 : Ferhat et al.
