## Supplementary material for "A peptide-neurotensin conjugate that crosses the blood-brain barrier induces pharmacological hypothermia associated with anticonvulsant, neuroprotective and anti-inflammatory properties following status epilepticus in mice": Table S5

Table S5. Mouse and Rat TaqMan probes used for qPCR analysis

| Gene name | Gene description | Probe ID |
| --- | --- | --- |
| *CIRBP* | *Cold Inducible RNA Binding Protein* | Mm00483336 |
| *CIRBP* | *Cold Inducible RNA Binding Protein* | Rn00579806 |
| *RBM3* | *RNA Binding Motif Protein 3* | Mm00812518 |
| *RBM3* | *RNA Binding Motif Protein 3* | Rn01525079 |
| *GAPDH* | *Glyceraldhyde-3-phosphate dehydrogenase* | Mm01253033 |
| *GAPDH* | *Glyceraldhyde-3-phosphate dehydrogenase* | Rn01253033 |

Table S5 : Ferhat et al.
